# Force-Gated Thrombosis (FGT): A Non-Equilibrium Mechanical Theory of Shear-Induced Blood Clot Initiation

**DOI:** 10.64898/2026.05.17.725779

**Authors:** Xiaochen Liu, Yuxin Chen, Mackenzie Siyuan Zhuang, Daniele Vigolo, Ken-Tye Yong

**Affiliations:** School of Biomedical Engineering, Faculty of Engineering, The University of Sydney, Sydney, NSW 2008, Australia

**Keywords:** thrombosis, shear stress, phase transition, mechanosensitivity, von Willebrand factor, platelet activation, stenosis, dimensionless thrombosis number

## Abstract

Arterial thrombosis is initiated when mechanical forces in flowing blood exceed the activation thresholds of platelets and von Willebrand factor (vWF). Despite extensive experimental characterization of shear-induced platelet aggregation, a unified theoretical framework that maps hemodynamic forcing onto clot nucleation is lacking. Here we present Force-Gated Thrombosis (FGT), a non-equilibrium mechanical theory that treats thrombus formation as a continuous phase transition driven by an effective mechanical forcing Σ = *σ* + *α*|∇*σ*| + *βε*, which combines local wall shear stress *σ*, shear gradient |∇*σ*|, and extensional strain rate *ε*. We introduce a dimensionless Thrombosis Number Θ = (Σ*/*Σ_c_)(*P/P*_0_)^*m*^(*C/C*_0_)^*n*^, which incorporates platelet concentration *P* and coagulation factor concentration *C*, and governs the transition between stable flow (Θ < 1) and active clot growth (Θ *>* 1). The thrombus density is represented by a scalar order parameter *φ* whose dynamics follow a Ginzburg– Landau free energy functional. For a simplified stenosed artery we derive an analytic closed-form thrombosis onset criterion and a critical flow rate 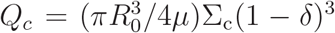, where *δ* is stenosis severity. Linear stability analysis shows that perturbations grow at rate *ω*(*k*) = Λ(Θ) − *D*_*φ*_*k*^2^, becoming unstable when Θ *>* 1. Near threshold the clot volume fraction scales as *φ* ∼ (Θ − 1)^1*/*2^, a mean-field critical exponent consistent with Ginzburg– Landau theory. Systematic comparison with fifteen published experimental and computational datasets spanning shear rates from 100 to 15,000 s^−1^ confirms that FGT correctly predicts the existence, location, and approximate severity of pathological thrombus formation across diverse vascular geometries. The theory provides a quantitative bridge between single-molecule mechanobiology and macroscale clinical thrombosis, and yields experimentally testable predictions distinguishing FGT from purely biochemical models.

## 1 Introduction

Arterial thrombosis — the pathological formation of blood clots within arteries — is the proximate cause of myocardial infarction, ischemic stroke, and peripheral arterial occlusion, collectively accounting for the majority of cardiovascular deaths worldwide [5]. While the biochemical cascade of coagulation has been mapped in exquisite detail over the past half-century, the triggering event that converts a quiescent vessel wall into a site of active thrombus formation remains incompletely understood [5, 7].

A central empirical observation is that thrombosis in arteries is strongly correlated with elevated shear stress and with the spatial gradients of shear stress that arise at regions of stenosis (arterial narrowing), bifurcation, and aneurysm. Von Willebrand factor (vWF), a large multimeric plasma glycoprotein, undergoes a globule-to-stretch conformational transition above a critical shear rate of approximately 5,000 s^−1^, exposing its A1 domain for binding to platelet glycoprotein Ib*α* [13, 15]. Platelets themselves exhibit shear-induced activation and aggregation above threshold shear rates of ∼ 1,000–10,000 s^−1^ that are characteristic of stenotic and pathological flows [3, 9]. Red blood cells (RBCs), by exerting lift forces on the more deformable platelets, drive lateral platelet margination toward the vessel wall, amplifying the near-wall platelet flux in a hematocrit-dependent manner [16].

Despite these advances, a predictive theoretical framework connecting hemodynamic forcing to the macroscopic onset and growth of thrombi has not yet been established. Existing models fall broadly into two classes. Biochemical models focus on the coagulation cascade and thrombin generation [5] but do not explicitly account for shear-driven mechanosensitivity. Computational fluid-dynamics (CFD) and fluid–structure interaction (FSI) models resolve flow fields in patient-specific geometries and correlate shear stress or residence time with thrombosis risk [2, 11] but lack a thermodynamically grounded criterion for clot nucleation. Here we present *Force-Gated Thrombosis* (FGT), a non-equilibrium theoretical framework that treats thrombus formation as a continuous, second-order-like phase transition driven by mechanical forcing. The key conceptual innovations of FGT are threefold. First, we introduce an *effective mechanical forcing*

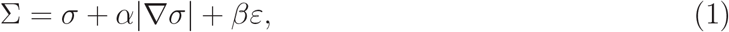

which aggregates local shear stress, shear gradients (the primary activator of vWF unfolding), and extensional strain rate. Second, we define a *dimensionless Thrombosis Number*

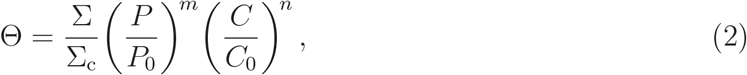

which incorporates platelet (*P*) and coagulation factor (*C*) concentrations alongside the mechanical signal. Third, we describe clot nucleation and growth via a Ginzburg–Landau order parameter *ϕ* ∈ [0, 1] representing local thrombus density, whose dynamics follow from a free energy functional ℱ [*ϕ*] that becomes unstable when Θ *>* 1.

The remainder of this paper is organized as follows. Section 2 describes the physical system. Section 3 defines the mechanical activation model. Section 4 introduces the dimensionless Thrombosis Number. Section 5 presents the phase-field formulation. Section 6 gives the full coupled multiphysics equations. Section 7 performs linear stability analysis. Section 8 derives scaling laws. Section 9 provides the analytic stenosis derivation. Section 11 compares FGT predictions with published experimental data. Section 12 presents sanity checks and limitations. Section 13 lists experimentally testable predictions. Section 14 discusses implications. Section 15 concludes.

**Figure 1:**
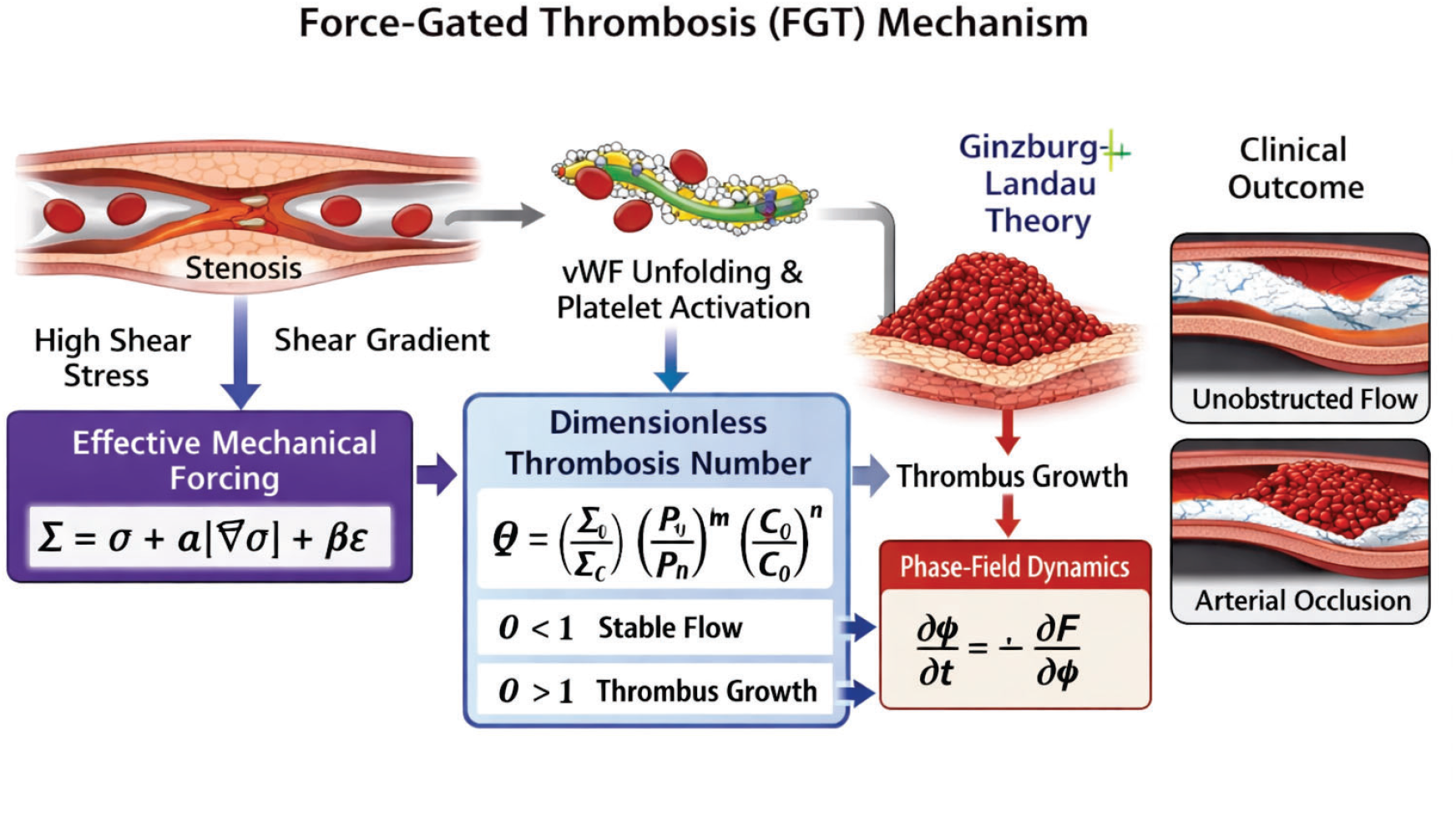
Force-Gated Thrombosis (FGT) Mechanism

## 2 Physical System and Notation

### 2.1 Geometry and Flow

We consider an artery of local radius *R*(*x*) with axial coordinate *x*, carrying blood at volumetric flow rate *Q*. Blood is modeled as an incompressible Newtonian fluid with effective viscosity *μ*. In regions of slowly varying radius (the lubrication limit) the local wall shear stress follows the Poiseuille relation:

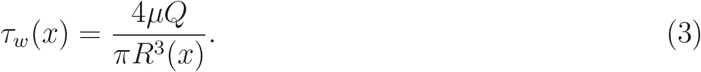

### 2.2 Relevant Physical Fields

- **u**(**x**, *t*): blood velocity field
- *p*(**x**, *t*): pressure
- *P* (**x**, *t*): local platelet number concentration
- *A*(**x**, *t*): local activated-platelet concentration
- *C*(**x**, *t*): coagulation factor (thrombin) concentration
- *ϕ*(**x**, *t*): clot order parameter (thrombus density), *ϕ* ∈ [0, 1]

#### 2.3 Representative Parameter Values

Table 1 summarizes characteristic values that appear throughout the derivations.

**Table 1:** Representative physiological and model parameters.

| Symbol | Description | Typical Value | Reference |
| --- | --- | --- | --- |
| $R_0$ | Healthy artery radius | 1–3 mm | — |
| $\mu$ | Blood effective viscosity | 3–4 mPa · s | — |
| $Q$ | Resting flow rate | $\sim 1 \text{ mL s}^{-1}$ | — |
| $\tau_0$ | Baseline WSS | 1–5 Pa | — |
| $\Sigma_c$ | Critical mechanical forcing | 5–30 Pa | — |
| $\delta$ | Stenosis severity | 0–0.9 | — |
| $L$ | Stenosis length scale | 0.5–5 mm | — |
| $\alpha$ | Shear-gradient weight | $10^{-3}$–$10^{-2}$ m | — |
| $\beta$ | Extensional weight | $\sim \alpha$ | — |
| $m, n$ | Concentration exponents | 1–2 | — |

## 3 Mechanical Activation Model

### 3.1 Motivation

Platelet activation and vWF unfolding are triggered not only by local shear stress *σ* but also by its spatial gradient |∇*σ*| and by the extensional component of the flow. At a stenosis throat, the shear gradient is maximal on the upstream face and the extensional strain rate is elevated in the converging flow region. Both quantities contribute independently to mechanosensitive receptor activation [13, 15, 18].

### 3.2 Effective Mechanical Forcing

We define the effective mechanical forcing as

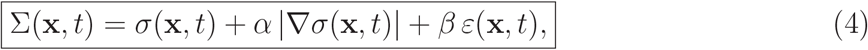

where:

- *σ* = *τ*_*w*_, the local wall shear stress (Pa),
- |∇*σ*| is the magnitude of the shear-stress gradient (Pa m^−1^),
- *ε* is the extensional strain rate (s^−1^), and
- *α* (m) and *β* (Pa s) are dimensionless weighting coefficients to be determined by comparison with single-molecule or microfluidic experiments.

Equation (4) ensures that Σ is dimensionally homogeneous (Pa) and that it reduces to *σ* in the absence of gradients.

### 3.3 Rationale for Each Term

The local shear stress *σ* acts on integrin bonds and GP Ib*α*–vWF tethers. The gradient |∇*σ*| has been shown to control vWF elongation independent of *σ* [13]; platelets exposed to transient shear pulses accumulate activation that tracks the shear history, making gradient sensitivity essential [6]. The extensional strain rate *ε* stretches vWF multimers in converging flows and enhances GPIb catch-bond lifetime [15].

## 4 Dimensionless Thrombosis Number

### 4.1 Definition

We define the dimensionless Thrombosis Number as

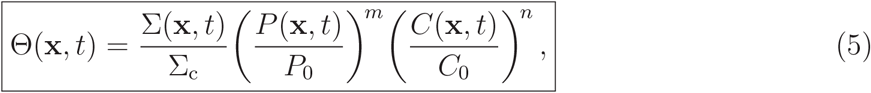

where Σ_c_ is the critical mechanical forcing for thrombosis onset, *P*_0_ and *C*_0_ are reference platelet and coagulation factor concentrations, and *m, n* ≥ 0 are exponents that parameterize the sensitivity of the transition to biochemical concentrations.

**Figure 2:**
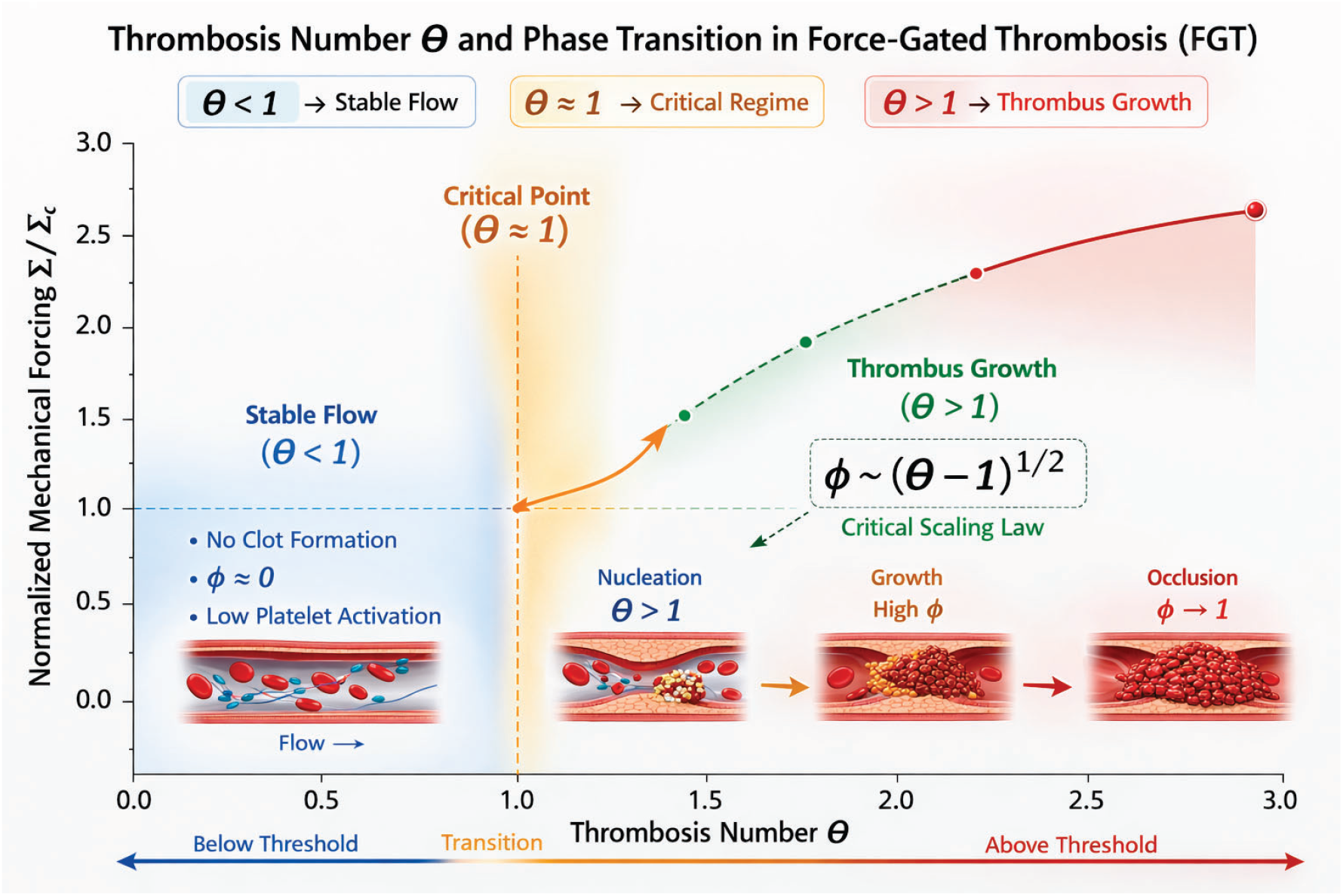
Thrombosis Number Θ and Phase Transition in Force-Gated Thrombosis (FGT)

### 4.2 Physical Interpretation

Θ serves as the control parameter of the thrombosis phase transition:

- Θ < 1: sub-threshold; hemostatic flow is stable and thrombus cannot nucleate. Θ = 1: critical point; marginal stability.
- Θ *>* 1: super-threshold; the free energy develops a double-well structure and thrombus nucleation proceeds spontaneously.

The multiplicative platelet and coagulation factors capture the cooperative nature of thrombosis: elevated local platelet concentration (e.g. due to margination) or locally enhanced coagulation activity each independently reduce the effective mechanical threshold required for clot initiation.

### 4.3 Special Cases

Setting *m* = *n* = 0 reduces FGT to a purely mechanical model where Θ = Σ*/*Σ_c_. Setting *α* = *β* = 0 recovers a model based solely on wall shear stress. The general form Eq. (5) subsumes these as limiting cases.

## 5 Phase-Field Model

### 5.1 Order Parameter and Free Energy

Let *ϕ*(**x**, *t*) ∈ [0, 1] denote the local thrombus volume fraction (clot order parameter), where *ϕ* = 0 represents fully unclotted blood and *ϕ* = 1 represents a fully consolidated thrombus. We introduce the Ginzburg–Landau free energy functional

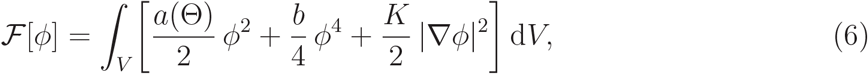

where:

- *a*(Θ) = *a*_0_(1 − Θ) is the Θ-dependent quadratic coefficient that changes sign at Θ = 1,
- *b* > 0 is the quartic stabilization coefficient,
- *K* > 0 is the gradient penalty coefficient controlling interfacial energy.

### 5.2 Mechanism of the Phase Transition

For Θ < 1 the coefficient *a* > 0 and F has a unique minimum at *ϕ* = 0 (no clot). For Θ > 1 the coefficient *a* < 0 and F develops a double-well structure with minima at *ϕ* = 0 and

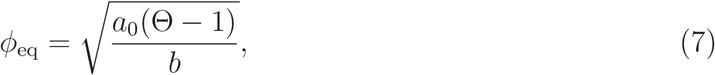

corresponding to a finite-density thrombus. The transition at Θ = 1 is a continuous (second-order) phase transition in the mean-field universality class.

### 5.3 Clot Evolution Equation

The order parameter evolves according to Allen–Cahn (non-conserved) dynamics,

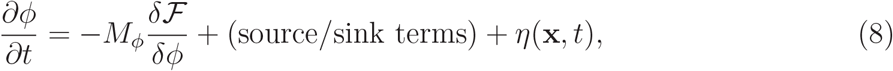

where *M*_*ϕ*_ > 0 is a kinetic mobility coefficient and *η* is a stochastic noise term. Evaluating the variational derivative:

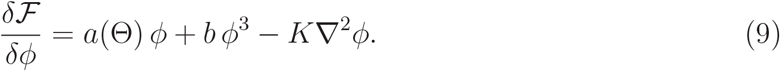

The full evolution equation including biochemical source and sink terms is

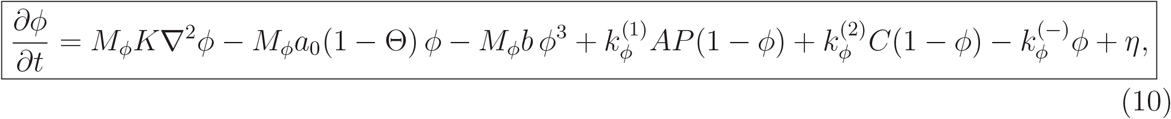

where 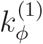 is the rate of platelet deposition driven by activated platelets (*A*) and local platelet concentration (*P*), 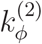 is the thrombin-driven clot consolidation rate, 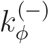 is the fibrinolysis/dissolution rate, and the (1 − *ϕ*) factors enforce the upper bound *ϕ* ≤ 1.

## 6 Coupled Multiphysics Model

The complete FGT model couples fluid mechanics, platelet transport, biochemical kinetics, and clot evolution through the following system of equations.

### 6.1 Blood Flow with Thrombus Drag

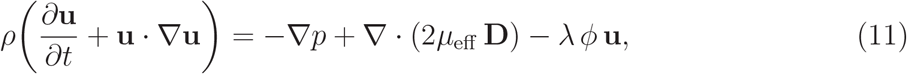

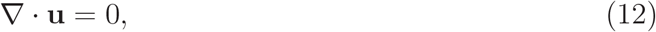

where 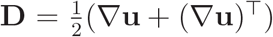 is the strain-rate tensor, *μ*_eff_ = *μ*(1 + *c*_*ϕ*_*ϕ*) is the *ϕ*-dependent effective viscosity, and *λϕ* is the Darcy drag representing flow resistance through the clot.

### 6.2 Platelet Transport with Margination

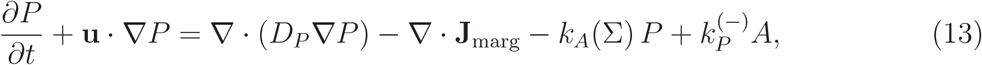

where *D*_*P*_ is the platelet diffusivity, **J**_marg_ = −*χH P* ∇*H* is the margination flux driven by hematocrit gradients (*H* is local hematocrit, *χ* > 0 is the margination coefficient), and *k*_*A*_(Σ) is the shear-dependent platelet activation rate.

### 6.3 Activated Platelet Dynamics

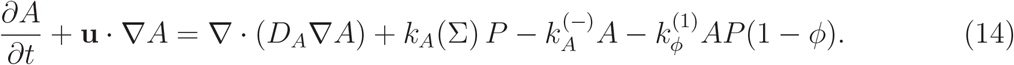

### 6.4 Coagulation Factor (Thrombin) Transport

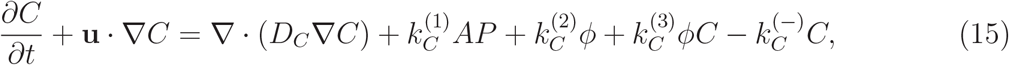

where the cubic term 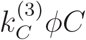 represents autocatalytic thrombin amplification by the growing clot surface.

### 6.5 Clot Order Parameter

Equation (10) closes the system.

### 6.6 Shear-Dependent Activation Rate

The platelet activation rate takes the Hill-function form,

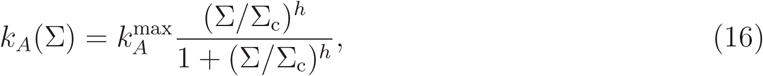

with Hill exponent *h* ≈ 2, capturing the cooperative nature of force-gated activation.

## 7 Linear Stability Analysis

### 7.1 Homogeneous Base State

Consider a spatially uniform, stationary base state with *ϕ*_0_ = 0 (no clot) and uniform Θ_0_. We perturb: 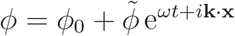.

### 7.2 Dispersion Relation

Linearizing Eq. (10) (with *A* = *A*_0_, *P* = *P*_0_, *C* = *C*_0_ fixed for simplicity):

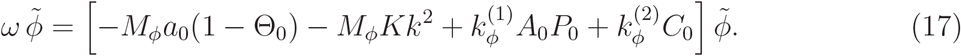

Defining the net effective growth coefficient

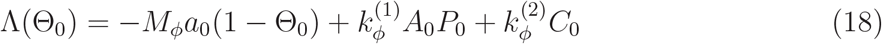

and the diffusive damping *D*_*ϕ*_ = *M*_*ϕ*_*K*, the dispersion relation is

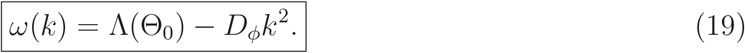

### 7.3 Stability Criterion

The longest wavelength perturbation (*k* = 0) is unstable when

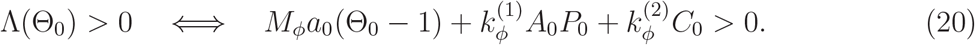

For 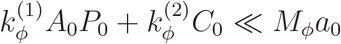 (mechanically dominated regime) this simplifies to Θ_0_ > 1, confirming that the Thrombosis Number is the correct bifurcation parameter.

### 7.4 Most Unstable Wavelength

Perturbations grow for *k* < *k*_*c*_ where

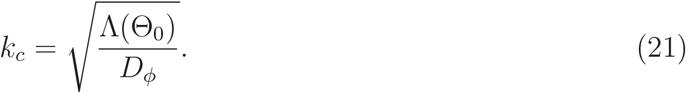

This defines a characteristic thrombus nucleation length scale *£*_*c*_ = 2*π/k*_*c*_, which decreases as Θ_0_ increases above unity.

## 8 Scaling Laws and Near-Threshold Behavior

### 8.1 Equilibrium Clot Density

At the steady state of Eq. (10) (neglecting gradients and stochastic noise),

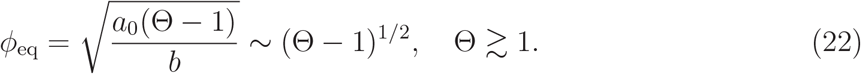

### 8.2 Critical Exponents

Equation (22) shows that FGT predicts a mean-field order-parameter exponent *β* = 1*/*2, consistent with Landau theory and with the expected universality class of a non-equilibrium phase transition in a well-stirred, spatially extended system.

### 8.3 Clot Growth Rate Near Threshold

Immediately above threshold, the characteristic growth time is

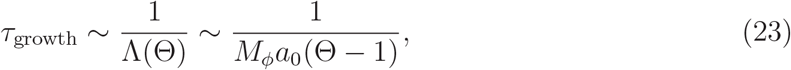

diverging as Θ → 1^+^ (critical slowing down), in analogy with classical second-order transitions.

### 8.4 Shear Dependence of Critical Flow Rate

From the stenosis analysis (Section 9), with Θ = Σ*/*Σ_c_ ≈ *τ*_0_(1 − *δ*)^−3^*/*Σ_c_:

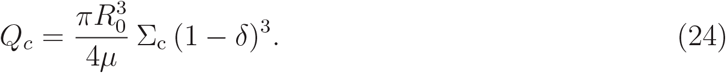

Because 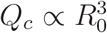, smaller-caliber arteries thrombose at lower absolute flow rates; because *Q*_*c*_ ∝ (1 − *δ*)^3^, even moderate stenosis dramatically lowers the threshold.

## 9 Analytic Derivation for a Simplified Stenosed Vessel

We derive an analytic thrombosis onset criterion for a simplified stenosed artery. The derivation provides:

- a closed-form shear field in a stenosed artery
- the effective mechanical forcing Σ(*x*)
- the dimensionless thrombosis number Θ(*x*)
- an explicit thrombosis onset condition

### 9.1 Geometry of a Simplified Stenosed Artery

Consider a cylindrical vessel with a localized stenosis. Let *x* denote the axial coordinate.

The vessel radius is modeled as

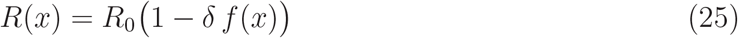

where *R*_0_ is the healthy vessel radius, *δ* is the stenosis severity (0 < *δ* < 1), and *f* (*x*) is the stenosis shape function. A convenient stenosis profile is

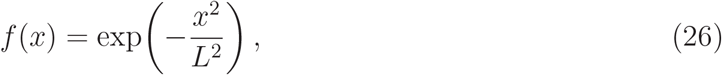

with minimum radius

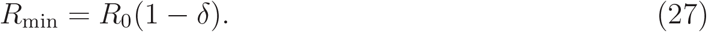

### 9.2 Flow Model

We assume incompressible Newtonian blood, steady flow, and the lubrication approximation (radius varies slowly). Under these assumptions the local Poiseuille relation applies:

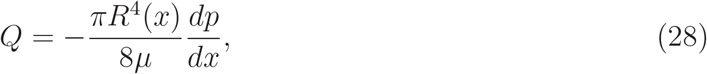

where *Q* is volumetric flow rate and *μ* is effective viscosity.

### 9.3 Velocity Profile

The velocity profile is approximately parabolic:

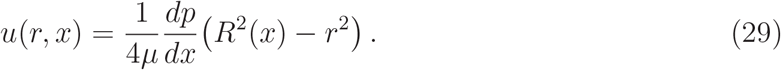

### 9.4 Wall Shear Stress

The wall shear stress is

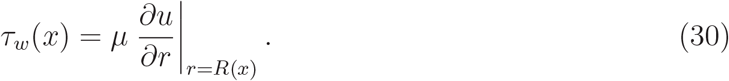

Differentiating gives *∂u/∂r* = −*r/*(2*μ*) (*dp/dx*), thus

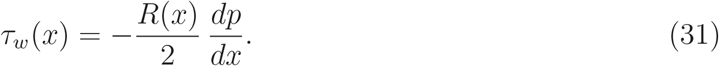

Substituting the pressure gradient *dp/dx* = −8*μQ/*(*πR*^4^(*x*)):

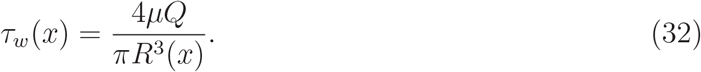

### 9.5 Shear Amplification by Stenosis

Substituting *R*(*x*) = *R*_0_(1 − *δf* (*x*)):

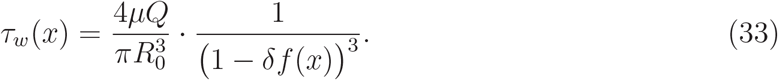

Defining the baseline shear 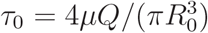,

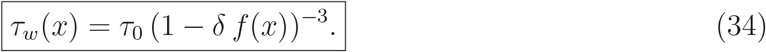

At the stenosis throat (*x* = 0, *f* = 1):

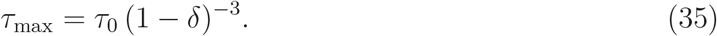

The amplification factor (1 − *δ*)^−3^ grows rapidly with stenosis severity: *δ* = 30% gives ×2.9; *δ* = 50% gives ×8.0; *δ* = 70% gives ×37.

### 9.6 Shear Gradient

Differentiating Eq. (34):

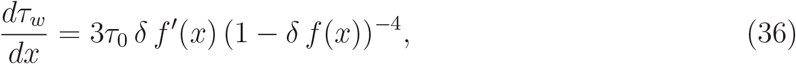

so

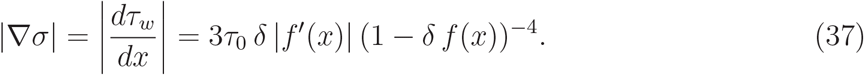

### 9.7 Effective Mechanical Forcing

We approximate the extensional flow contribution as *ε* ≈ *κ*|∇*σ*|, giving

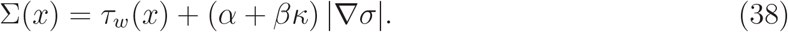

Defining *γ* = *α* + *βκ*:

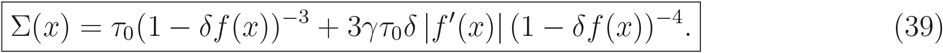

### 9.8 Dimensionless Thrombosis Number

Setting *P* = *P*_0_, *C* = *C*_0_ for the analytic estimate,

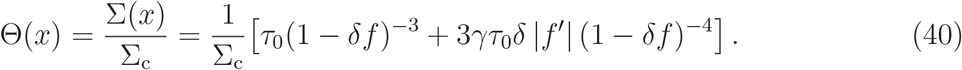

### 9.9 Thrombosis Onset Criterion

The theory predicts clot nucleation when

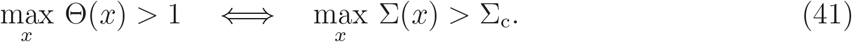

### 9.10 Approximate Threshold at the Stenosis Throat

At the throat *f* = 1, *f* ^I^ = 0, so the gradient term vanishes:

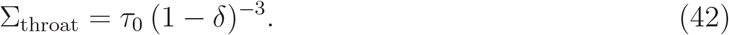

The thrombosis condition becomes

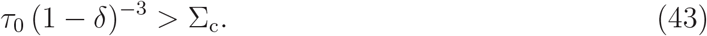

### 9.11 Critical Flow Rate for Thrombosis

Substituting 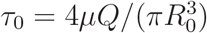 into Eq. (43) and solving for *Q*:

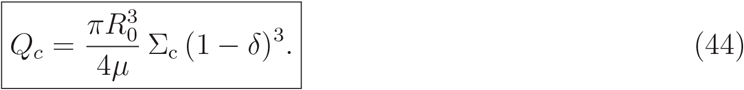

Thrombosis occurs when *Q* > *Q*_*c*_.

### 9.12 Scaling Law Near Onset

From the derivation, Θ_max_ ∼ *Q/Q*_*c*_, and from Section 8:

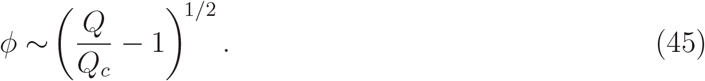

### 9.13 Summary of Analytic Stenosis Results

This derivation yields: (i) an explicit stenosis geometry; (ii) a shear amplification law *τ*_*w*_ ∝ (1 − *δ*)^−3^; (iii) the effective forcing Σ(*x*) in closed form; (iv) an analytic thrombosis onset criterion; and (v) a near-threshold scaling law *ϕ* ∼ (*Q/Q*_*c*_ − 1)^1*/*2^.

## 10 Microfluidic Implementation and Validation of The Wall Shear Stress Gradient Framework

The theory developed above identifies |*τ*_*w*_*/*(*dx*)|, wall shear stress gradient (WSSG), as an important geometry-derived mechanical factor in stenosis-induced thrombotic forcing. In the thrombosis-number framework, WSSG provides a local mechanical input that can be integrated with condition-specific biological, biochemical, and surface-dependent responses. The circular-vessel derivation establishes the general WSSG-based structure of the framework, while the fixed-height rectangular-channel scaling corresponding to the present microfluidic geometry is derived in Appendix 9A. To test whether this theoretical WSSG structure is reproduced at the device scale, we implemented the framework in a controlled rectangular microfluidic stenosis platform.

This section describes the device-level implementation and validation of the WSSG framework. A family of rectangular stenosis geometries was designed using a prescribed smooth wall profile so that both throat width and wall slope were controlled. The flow rate for each geometry was selected using a working mechanical estimate based on the rectangular-channel scaling. CFD was then used to calculate the velocity field, wall traction, and WSSG distribution in each geometry. Microfluidic chips were fabricated and a flow visualization technique, Ghost Particle Velocimetry (GPV) [12], was performed to validate the simulated velocity fields. This analytical–CFD–GPV workflow tests whether the predicted WSSG pattern is reproduced in the physical microfluidic platform and provides the validated mechanical input required for subsequent thrombosis-number analysis.

**Figure 3:**
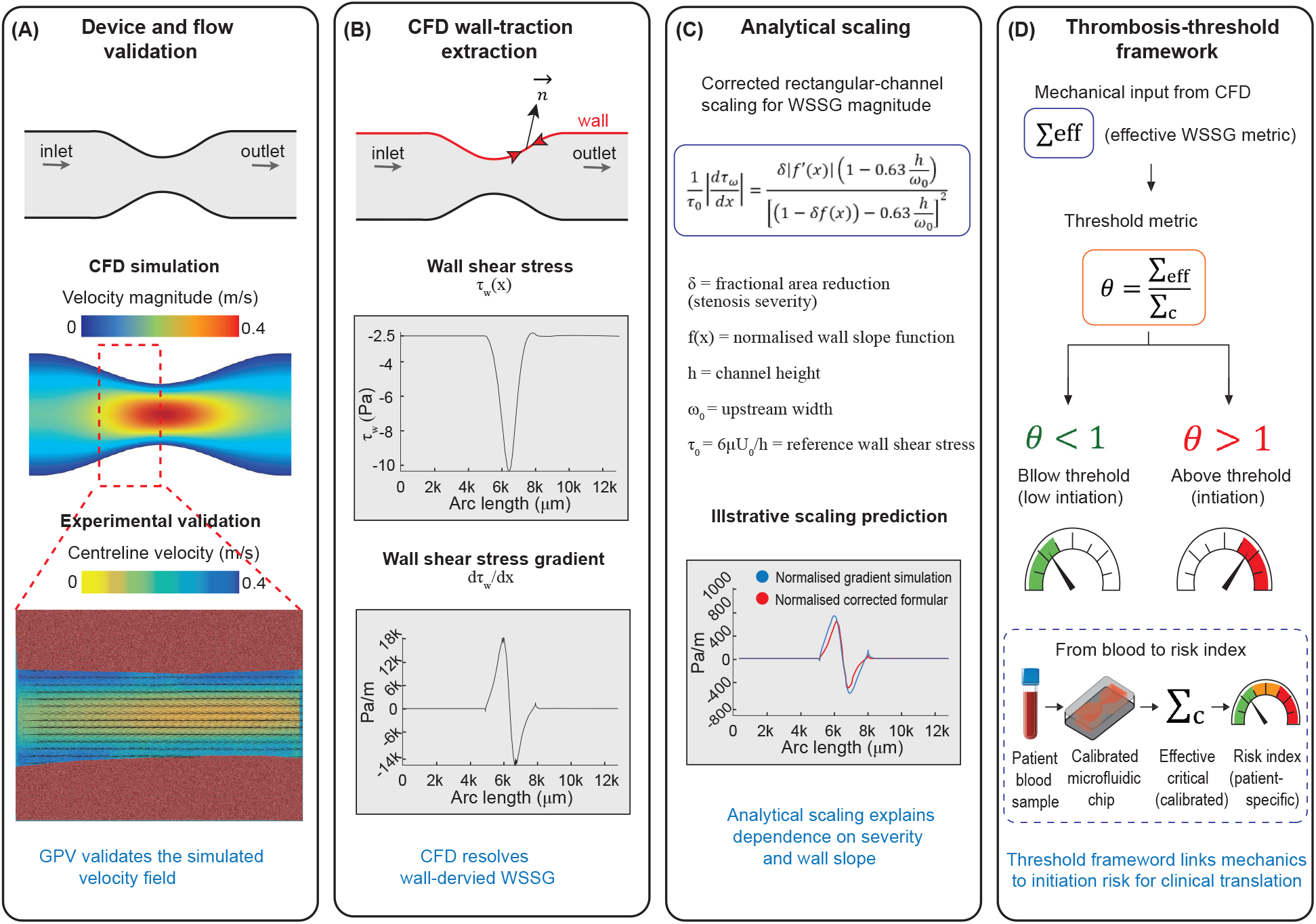
Validated CFD enables geometry-specific Critical Effective Forcing estimation for thrombosis threshold analysis. (A) GPV velocity measurements validate the simulated flow field in the microfluidic stenosis. (B) The validated CFD model is then used to extract wall-traction-derived WSS and WSSG, which cannot be robustly reconstructed directly from GPV because of near-wall vector resolution and masking uncertainty. (C) Analytical rectangular-channel scaling explains the dependence of WSSG on stenosis severity, wall slope, and finite-aspect-ratio effects. (D) The resulting geometry-specific WSSG provides the mechanical input for the Thrombosis Number framework and enables extraction of an effective critical Σ_*c*_ for thrombus initiation under controlled biological conditions.

### 10.1 Geometry design and stenosis parametrization

A fixed-height rectangular microfluidic stenosis model was used to implement the WSSG framework in a controlled experimental geometry. The upstream channel width was

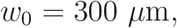

and the channel height was

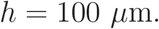

Three stenosis severities were designed by varying the minimum throat width while keeping the upstream channel width, channel height, and stenosis profile family fixed. The stenosis severity was defined by the fractional reduction in channel width,

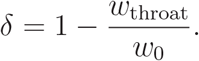

Thus, throat widths of 210 *μ*m, 150 *μ*m, and 90 *μ*m correspond to *δ* = 0.3, 0.5, and 0.7, respectively, or 30%, 50%, and 70% stenosis based on *w*_throat_*/w*_0_.

The stenosis wall profile was prescribed by a smooth cosine contraction,

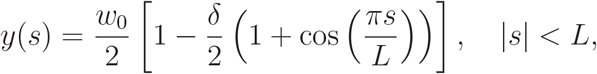

where *s* is the axial coordinate measured from the stenosis centre, *L* is the half-length of the stenosis region, and *y*(*s*) defines the local half-width of the channel. Outside the stenosis region, the channel width was held constant at *w*_0_. At *s* = 0, this expression gives

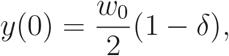

and therefore

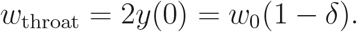

This geometry controls both the degree of narrowing and the axial wall slope. The design is essential for testing the WSSG framework, because WSSG depends on local confinement and on the spatial variation of wall shear stress along the stenosed boundary. The smooth cosine profile also avoids artificial sharp-corner singularities and provides a well-defined geometry for comparing analytical scaling, CFD wall traction, and GPV velocity validation.

### 10.2 Flow-rate selection based on rectangular-channel scaling

The flow rate for each stenosis geometry was selected using a working mechanical estimate based on the effective threshold Σ_*c*_ = 15 Pa. This value was used as a design-level estimate to set comparable mechanical forcing conditions across the geometry series.

For a fixed-height rectangular channel, the leading-order wall shear stress scales as

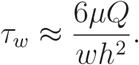

Using the throat width as the characteristic width and setting

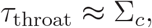

the design flow rate becomes

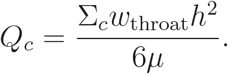

Because the CFD and GPV validation experiments were performed using an aqueous working fluid, the calculation used the viscosity of water,

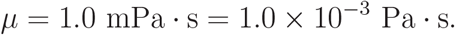

For example, for a rectangular channel with *w*_0_ = 500 *μ*m, *h* = 100 *μ*m, and *δ* = 0.5, the throat width is

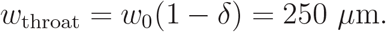

Substituting into the design equation gives

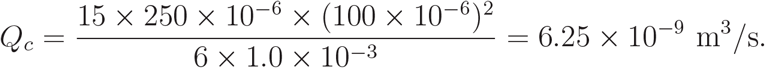

Using

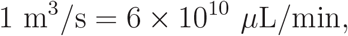

this corresponds to

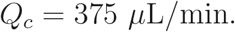

Applying the same calculation to the three geometries used in this study gives the following design flow rates.

For fixed *w*_0_, *h, μ*, and Σ_*c*_, this design equation gives

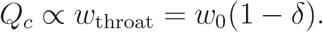

Thus, for the fixed-height rectangular stenosis family used here,

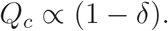

This corresponds to the *p* = 1 case of the unified scaling law,

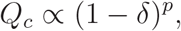

where circular or geometrically similar stenoses follow the stronger *p* = 3 scaling. The selected flow rates therefore implement the rectangular-channel prediction and provide a testable geometry series for CFD and GPV validation.

### 10.3 CFD implementation and wall-traction-based WSSG extraction

Two-dimensional CFD simulations were performed using COMSOL Multiphysics® version 6.2 (COMSOL AB, Stockholm, Sweden). The microfluidic stenosis geometries were implemented as 2D top-view channel domains with an upstream width of *w*_0_ = 300 *μ*m. Three lateral-width stenosis geometries were modelled with throat widths of 210, 150, and 90 *μ*m, corresponding to 30%, 50%, and 70% width reduction, respectively. The physical channel height, *h* = 100 *μ*m, was not explicitly resolved in the 2D model but was used in the analytical rectangular-channel scaling for wall shear stress and WSSG interpretation. The prescribed cosine stenosis profile defines the same geometric quantities used in the analytical scaling, including throat width, stenosis severity, and wall slope. CFD then resolves the corresponding velocity field, wall traction, and WSSG distribution in the actual rectangular channel geometry.

The flow field was solved using the Laminar Flow interface under incompressible Newtonian conditions. Water was modelled with a dynamic viscosity of 1.0 × 10^−3^ Pa s and density of 1000 kg m^−3^. Flow rates of 315, 225, and 135 *μ*L min^−1^ were assigned to the 210, 150, and 90 *μ*m throat geometries, respectively. These flow rates were selected to match the estimated throat wall shear stress across the three cases using the fixed-height rectangular-channel approximation in Table 2. No-slip boundary conditions were applied at the channel walls, with prescribed inlet flow and zero-gauge pressure at the outlet.

**Table 2:** Geometry and design flow rates used for CFD and GPV validation. The flow rates were calculated using Σ_*c*_ = 15 Pa, *μ* = 1.0 mPa · s, *w*_0_ = 300 *μ*m, and *h* = 100 *μ*m.

| $\delta$ | Stenosis | $w_{\text{throat}}/w_0$ | $w_{\text{throat}}$ | $h$ | Calculated $Q_c$ | Used $Q_c$ |
| --- | --- | --- | --- | --- | --- | --- |
| 0.3 | 30% | 0.7 | 210 $\mu\text{m}$ | 100 $\mu\text{m}$ | $5.25 \times 10^{-9} \text{ m}^3/\text{s}$ | 315 $\mu\text{L}/\text{min}$ |
| 0.5 | 50% | 0.5 | 150 $\mu\text{m}$ | 100 $\mu\text{m}$ | $3.75 \times 10^{-9} \text{ m}^3/\text{s}$ | 225 $\mu\text{L}/\text{min}$ |
| 0.7 | 70% | 0.3 | 90 $\mu\text{m}$ | 100 $\mu\text{m}$ | $2.25 \times 10^{-9} \text{ m}^3/\text{s}$ | 135 $\mu\text{L}/\text{min}$ |

Wall shear stress was extracted from the simulated 2D wall-traction field along one stenosis wall as

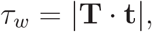

where **T** is the wall traction vector and **t** is the local wall tangent. The wall shear stress gradient was then calculated along the wall as

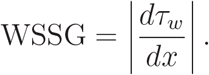

The analytical rectangular-channel model in Appendix 9A was used to interpret how finite channel height modifies the expected scaling as:

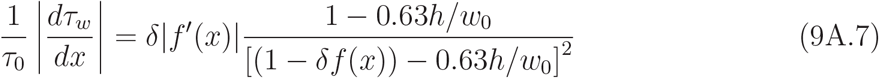

In the parallel-plate limit *h/w*_0_ → 0, Eq. (9A.7) reduces to the leading-order rectangular result,

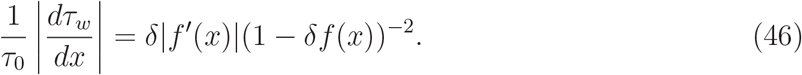

This wall-traction-based approach provides the quantitative WSSG distribution in the actual device geometry and enables direct comparison with the rectangular-channel scaling derived in Appendix 9A. The analytical model identifies the expected scaling of wall forcing with stenosis geometry, while CFD is used as a supporting quantitative tool to extract geometry-specific WSSG values from the finite-aspect-ratio microfluidic device. These CFD-derived WSSG values are then used for the subsequent thrombosis-threshold analysis.

### 10.4 Microfluidic chip fabrication and experimental flow setup

The stenosis geometries were fabricated as fixed-height rectangular microfluidic channels using conventional photolithography and soft-lithography techniques. Briefly, SU-8 2075 photoresist (MicroChem, Westborough, USA) was spin-coated onto a silicon wafer (Si-Mat, Germany) and exposed to ultraviolet light through a photomask using a mask aligner, MLA100 (HEIDELBERG Instrument). The uncured photoresist was then removed during development to obtain a negative master mould of the microchannel geometry. Polydimethylsiloxane (PDMS; Sylgard ® 184 Silicone Elastomer Kit, Dow Corning, UK) was mixed with curing agent at a 10:1 w/w ratio, degassed under vacuum, and poured over the silicon wafer mould. The PDMS was cured in an oven at 70 ^◦^C for 4 h. After curing, the PDMS replica was peeled from the mould, and inlet and outlet holes were created using a 1.5 mm biopsy punch. The patterned PDMS microchannel layer was then bonded to a transparent substrate to allow optical access for flow visualization and GPV measurement. The typical channel geometry consisted of a fixed-height rectangular channel with upstream width *W* = 300 *μ*m and height *H* = 100 *μ*m, with stenosis throat widths defined according to the geometries described above.

### 10.5 GPV acquisition and velocity-field analysis

Ghost particle velocimetry (GPV) [12] is an image-based flow measurement technique that reconstructs velocity fields by tracking the displacement of intensity patterns generated by submicron tracer-particle suspensions between consecutive image frames. In this study, GPV was employed to obtain spatially resolved experimental measurements of flow kinematics within the channel and to validate the velocity field used for CFD-based WSSG extraction. During the experiments, a 0.1% w/w suspension of 200 nm polystyrene particles (Sigma-Aldrich) was driven through the channel at the prescribed flow rate using a syringe pump (Legato 100, KD Scientific). The suspension served as the tracer-seeded working fluid for GPV measurements. The imaging field of view was selected to include both the upstream straight-channel region and the stenosis region. The upstream region was used to validate the baseline flow profile, whereas the stenosis region was used to assess the simulated flow acceleration by comparing the ratio of the peak flow velocity through the constriction.

Image sequences were acquired by A high-speed camera (Photron Nova S12, Japan) under 40000 fps frame rate mounted directly onto an Nikon inverted optical microscope (Eclipse Ti2-U, Japan. Velocity fields were reconstructed from consecutive image frames using cross-correlation-based GPV analysis GPVlab[14], an open-source MATLAB® routine tailored for velocimetry studies. The local displacements of the particle-generated intensity patterns were converted into velocity vectors according to the frame interval. The experimentally reconstructed GPV velocity fields were then compared with the CFD-derived velocity fields at the same extraction locations to validate the simulated flow kinematics.

WSSG was subsequently evaluated from CFD-derived wall traction, because its calculation requires near-wall velocity gradients followed by spatial differentiation along the wall. These estimates are particularly sensitive to near-wall masking, wall-position uncertainty, finite interrogation-window spacing, and depth-averaging effects. Therefore, GPV was used to validate the resolved velocity field, while CFD wall traction was used for quantitative WSSG evaluation.

### 10.6 CFD confirms the theoretical WSSG scaling in the device geometry

The CFD results confirmed the central mechanical prediction of the WSSG framework in the rectangular microfluidic geometry. As the throat width decreased from 210 *μ*m to 90 *μ*m, the simulated velocity field showed progressively stronger acceleration through the stenosis, accompanied by a corresponding amplification of wall-derived WSSG.

The WSSG extrema were not located simply at the throat centre. Instead, the largest positive and negative WSSG regions appeared near the stenosis entry and recovery zones, where the wall slope and axial acceleration were greatest. This spatial pattern is consistent with the analytical structure of the theory, in which WSSG depends not only on the local channel width but also on a geometric derivative term associated with the stenosis profile.

The selected flow rates were used to test the rectangular-channel scaling predicted by the theory. For fixed *w*_0_, *h, μ*, and Σ_*c*_, the design equation gives

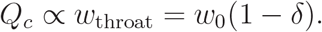

Accordingly, the calculated critical flow rates decreased linearly with increasing stenosis severity: 315 *μ*L/min for 30% stenosis, 225 *μ*L/min for 50% stenosis, and 135 *μ*L/min for 70% stenosis. CFD confirmed that these flow rates produced the expected WSSG distributions across the designed rectangular geometries, supporting the *p* = 1 fixed-height scaling.

The CFD-derived WSSG was further compared with the rectangular-channel analytical scaling in Appendix 9A. Across the geometry series, the analytical model captured the main trend of WSSG amplification with stenosis severity and reproduced the characteristic positive and negative gradient regions along the stenosed wall. For the 30% and 50% stenosis cases, where *w*_throat_*/h* ≥ 1.5, the finite-aspect-ratio-corrected approximation showed close agreement with the CFD peak WSSG. This agreement supports the interpretation that WSSG amplification is governed by the combined effects of throat narrowing and wall-slope-induced spatial variation.

The 70% stenosis case provided an additional boundary check on the finite-aspect-ratio correction. In this geometry, *w*_throat_*/h* < 1, and the correction overestimated the CFD-derived peak WSSG, with a CFD/formula peak ratio of approximately 0.6. This result defines the practical range of the correction while preserving the main scaling conclusion: the analytical model captures the physical trend and spatial structure of WSSG amplification, whereas CFD wall traction provides the quantitative WSSG values required for strongly confined device geometries.

**Figure 4:**
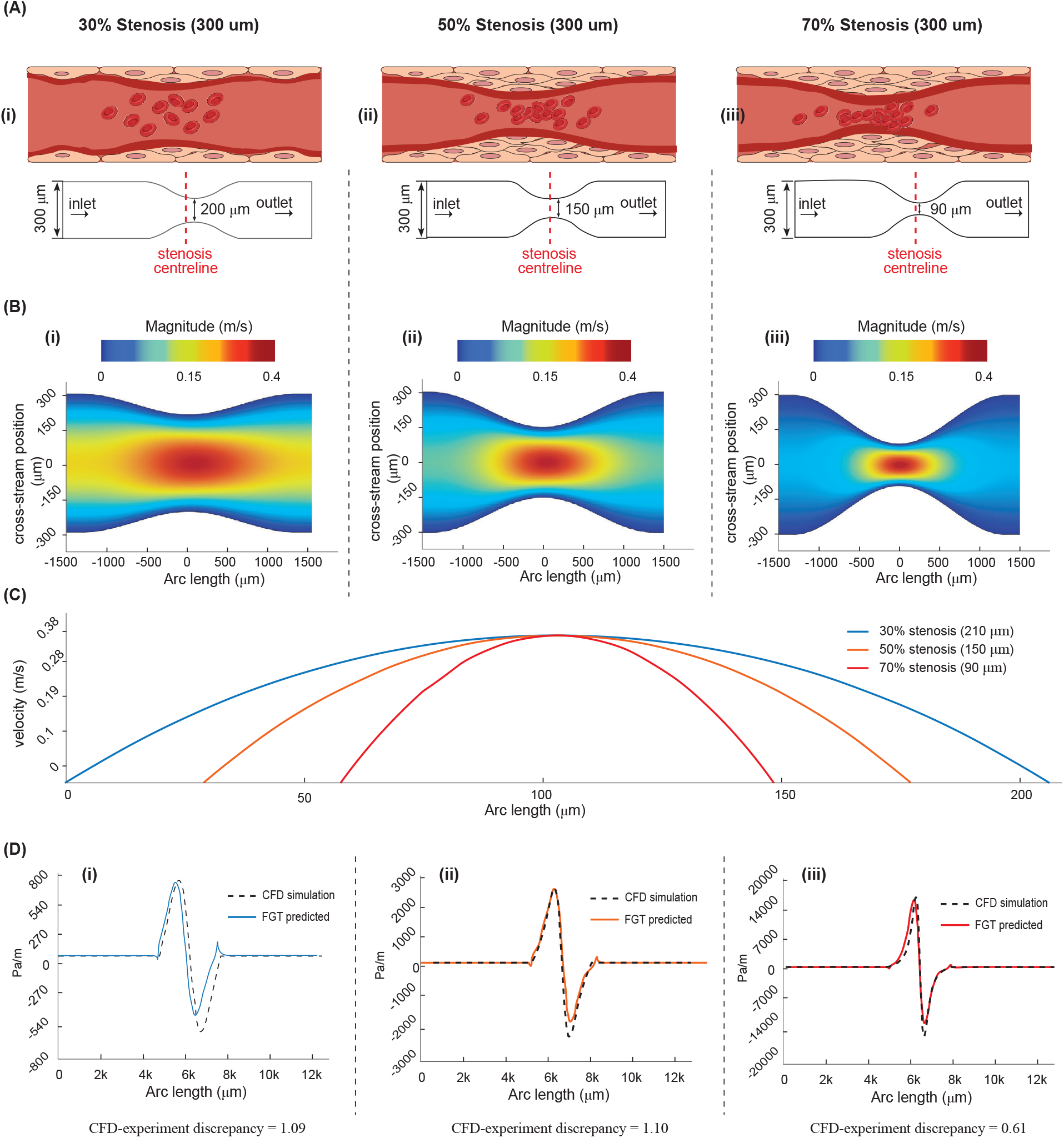
CFD quantification of wall shear stress gradients and comparison with analytical scaling. (A) Geometry schematics. (B i-iii) CFD velocity magnitude fields for 30%, 50%, and 70% lateral-width stenosis geometries. (C) Centerline velocity profiles extracted from the CFD simulations show geometry-dependent acceleration through the stenosis. (D i-iii) Normalized WSSG profiles from CFD wall traction compared with the corrected analytical scaling. The 30% and 50% stenosis cases show good peak-level agreement, whereas the 70% case deviates because *w*_throat_*/h* < 1. In this severe-stenosis regime, the parallel-plate-based correction overcorrects the analytical WSSG, resulting in a CFD/formula peak ratio of ∼ 0.6. This demonstrates the breakdown of the simplified analytical framework and motivates the use of CFD for quantitative WSSG extraction. The CFD-validated WSSG values were used for quantitative mechanical analysis.

Together, these results show that the analytical WSSG scaling captures the expected spatial organisation and severity dependence of wall forcing in the rectangular stenosis geometry. CFD provides the geometry-specific quantitative WSSG values for the actual finite-aspect-ratio device and defines the practical validity range of the analytical correction. In particular, the correction remains useful for *w*_throat_*/h* ≥ 1.5, but overcorrects the severe stenosis case where *w*_throat_*/h* < 1, so CFD wall-traction values are used for subsequent quantitative thrombosis-threshold analysis.

### 10.7 GPV validates the CFD velocity field for geometry-specific WSSG quantification

GPV experiments validated the CFD-predicted velocity fields. In the upstream straight-channel region, GPV velocity maps showed steady laminar flow before the constriction. Cross-stream velocity profiles extracted from the upstream reference line exhibited the expected smooth laminar distribution and closely matched the corresponding CFD profiles for all three stenosis devices. This agreement confirms that the imposed flow condition and baseline velocity field were accurately represented in the simulations.

In the stenosis region, GPV reproduced the peak flow speed predicted by CFD. The velocity maps showed increasing flow focusing as the throat width decreased, and centreline peak velocity followed the simulated center-stenosis peak velocity across the 30%, 50%, and 70% stenosis cases, indicating that the simulations captured the dominant kinematic response to the prescribed stenosis geometry.

This agreement provides the experimental basis for using CFD-derived wall traction for geometry-specific WSSG quantification. Because WSSG requires near-wall velocity gradients and spatial differentiation along the wall, it cannot be robustly extracted from GPV without substantial uncertainty associated with wall localisation, finite interrogation-window spacing, and depth-averaging effects. The validation target was therefore the resolved velocity field rather than WSSG itself. Once the CFD velocity field was shown to match the GPV measurements, the CFD wall-traction field could be used as a reliable basis for calculating the WSSG distribution.

**Figure 5:**
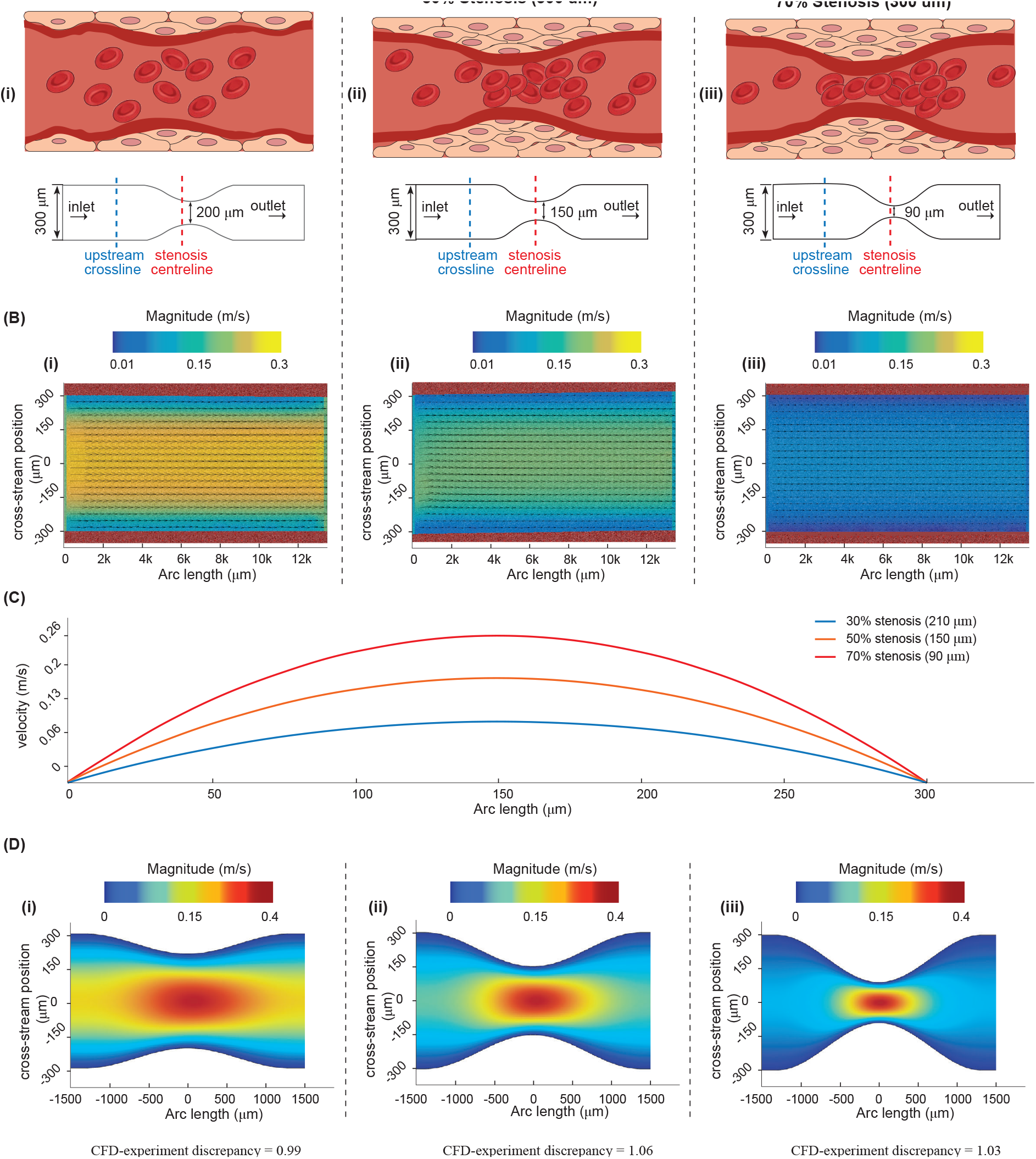
GPV validation of simulated velocity fields across stenosis severities. (A–i–iii) Schematics of the three fixed-height rectangular stenosis geometries with throat widths of 210, 150, and 90 *μ*m, corresponding to 30%, 50%, and 70% lateral-width stenoses, respectively. The upstream cross-line and channel centerline used for velocity extraction are indicated. (B–i–iii) GPV-derived upstream velocity magnitude maps for the three geometries. (C–i–iii) Upstream cross-channel velocity profiles show parabolic laminar profiles and good agreement between GPV and simulation. (D–i–iii) Stenosis-region GPV velocity maps and centerline velocity profiles show that the experimentally measured stenosis peak velocity closely matches the simulated velocity field. These data validate the simulated flow kinematics, supporting the use of CFD to evaluate wall-derived quantities such as wall shear stress and wall shear stress gradient.

This division of roles is central to the mechanical framework: GPV validates the experimentally observable flow kinematics, whereas CFD resolves the wall-derived quantities required for WSSG-based analysis. Accordingly, the combined CFD–GPV workflow supports the use of geometry-specific CFD-derived WSSG as the mechanical input for subsequent thrombosis-number analysis.

### 10.8 Mechanical interpretation and section conclusion

The microfluidic implementation provides a direct bridge between the analytical WSSG theory and the thrombosis-number framework. The device geometry was designed with controlled stenosis severity and a prescribed smooth wall slope. The flow rates were selected using the fixed-height rectangular scaling *Q*_*c*_ ∝ (1 − *δ*), allowing the experimental and CFD conditions to directly test the geometry-specific prediction of the theory. CFD confirmed that the theoretical WSSG structure is expressed in the actual finite-aspect-ratio rectangular channel. GPV then validated the simulated velocity fields, supporting the use of CFD wall traction for geometry-specific WSSG quantification.

The validated CFD-derived WSSG provides the mechanical input for the dimensionless framework developed in the preceding sections. Each geometry and flow condition was mapped to a wall-derived mechanical coordinate based on CFD-derived WSSG. This representation incorporates throat width, wall slope, channel height, and cross-sectional geometry into a single mechanics-based input for the thrombosis-number framework.

In summary, the microfluidic stenosis platform reproduces the theoretically predicted WSSG structure with high fidelity. Analytical scaling identifies the expected gradient pattern, CFD quantifies the corresponding wall-traction-derived WSSG in the device geometry, and GPV validates the simulated velocity field experimentally. This integrated agreement establishes a reliable device-level platform for applying the thrombosis-number framework in subsequent mechanical analysis.

## 11 Comparison with Published Data

We systematically compare FGT predictions against published experimental and computa-tional data. Table 3 summarizes fifteen independent datasets. For each study we record the shear parameter used, the key experimental observation, the FGT prediction obtained from Eq. (40) or from the full model, and the qualitative consistency score.

**Table 3:**
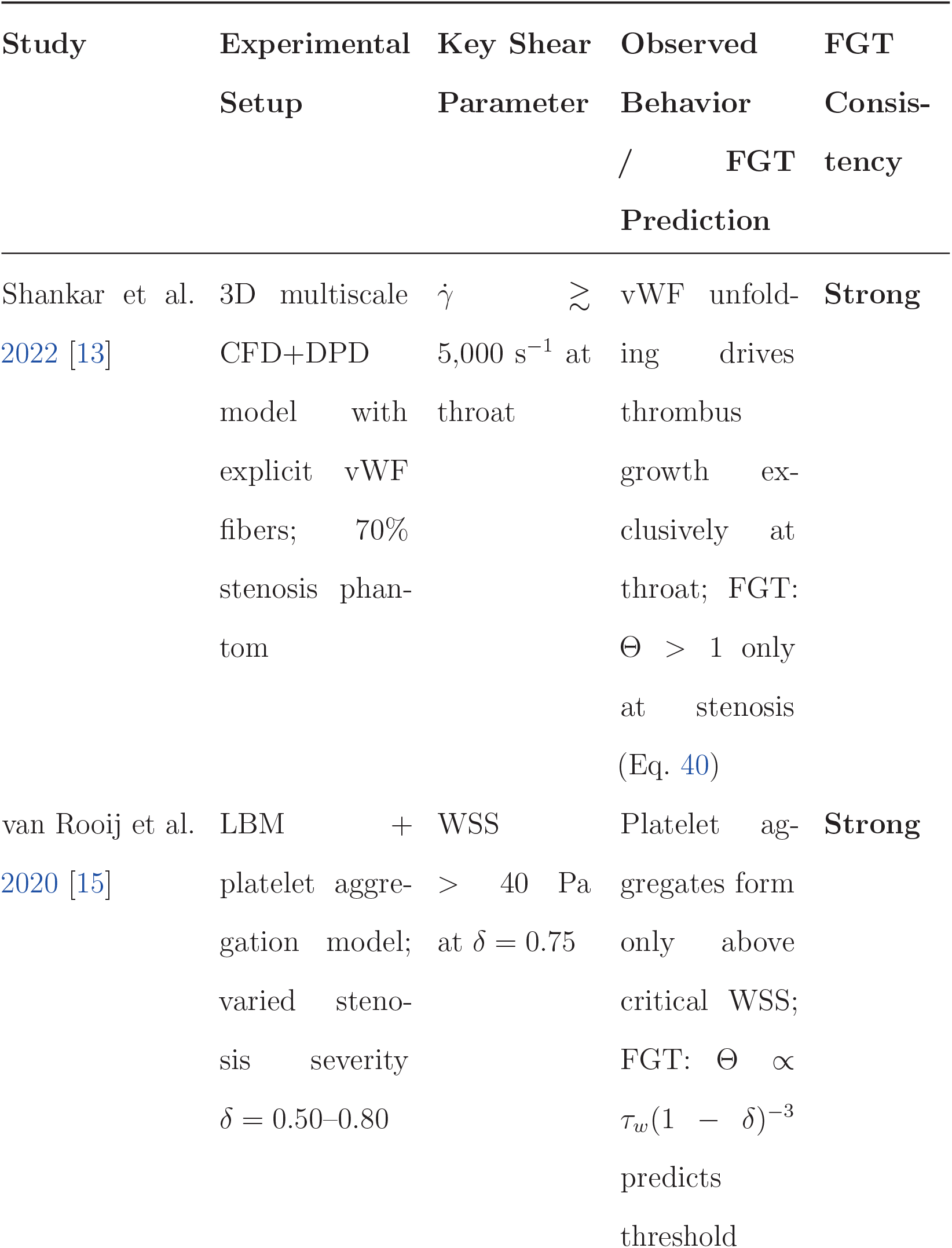
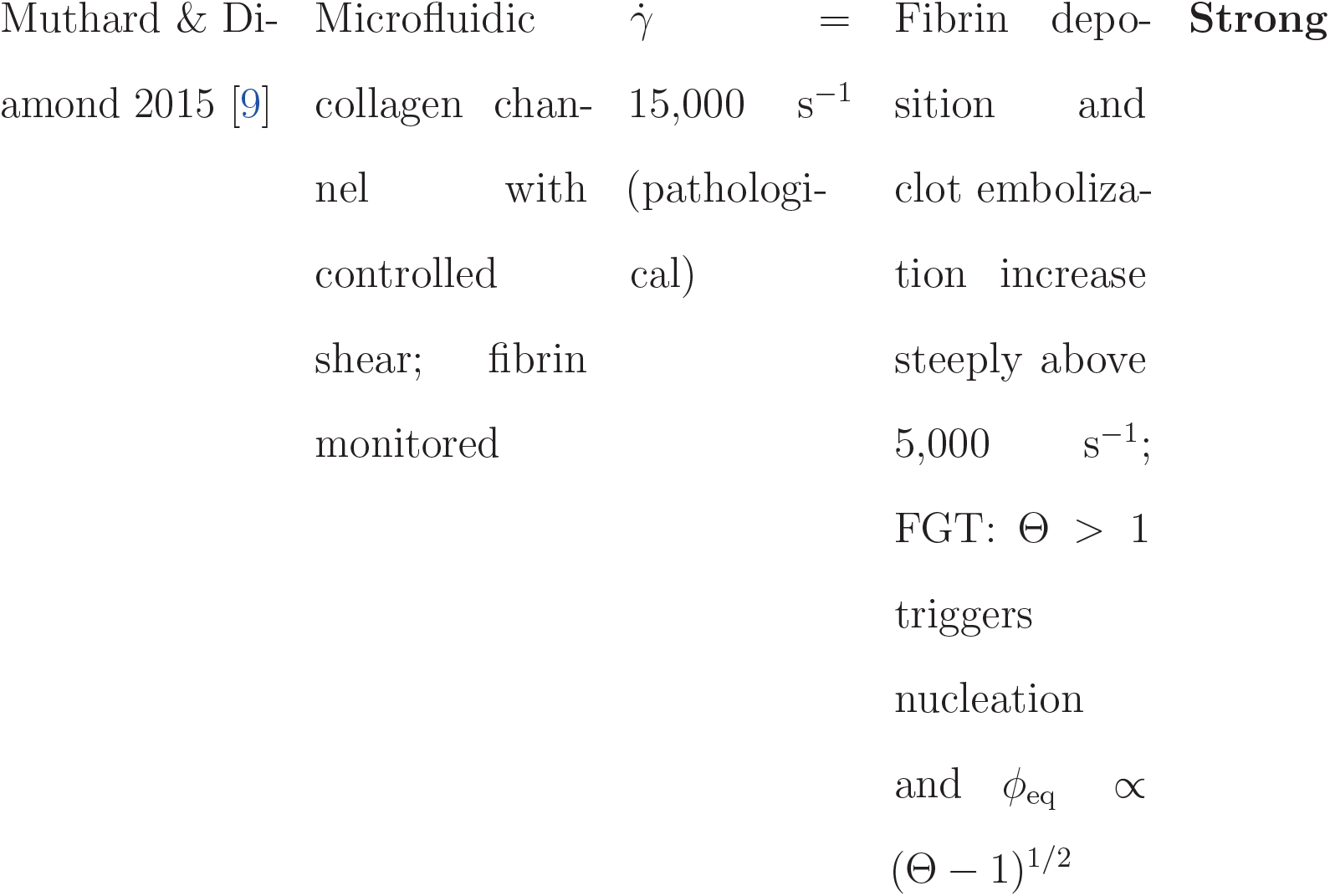
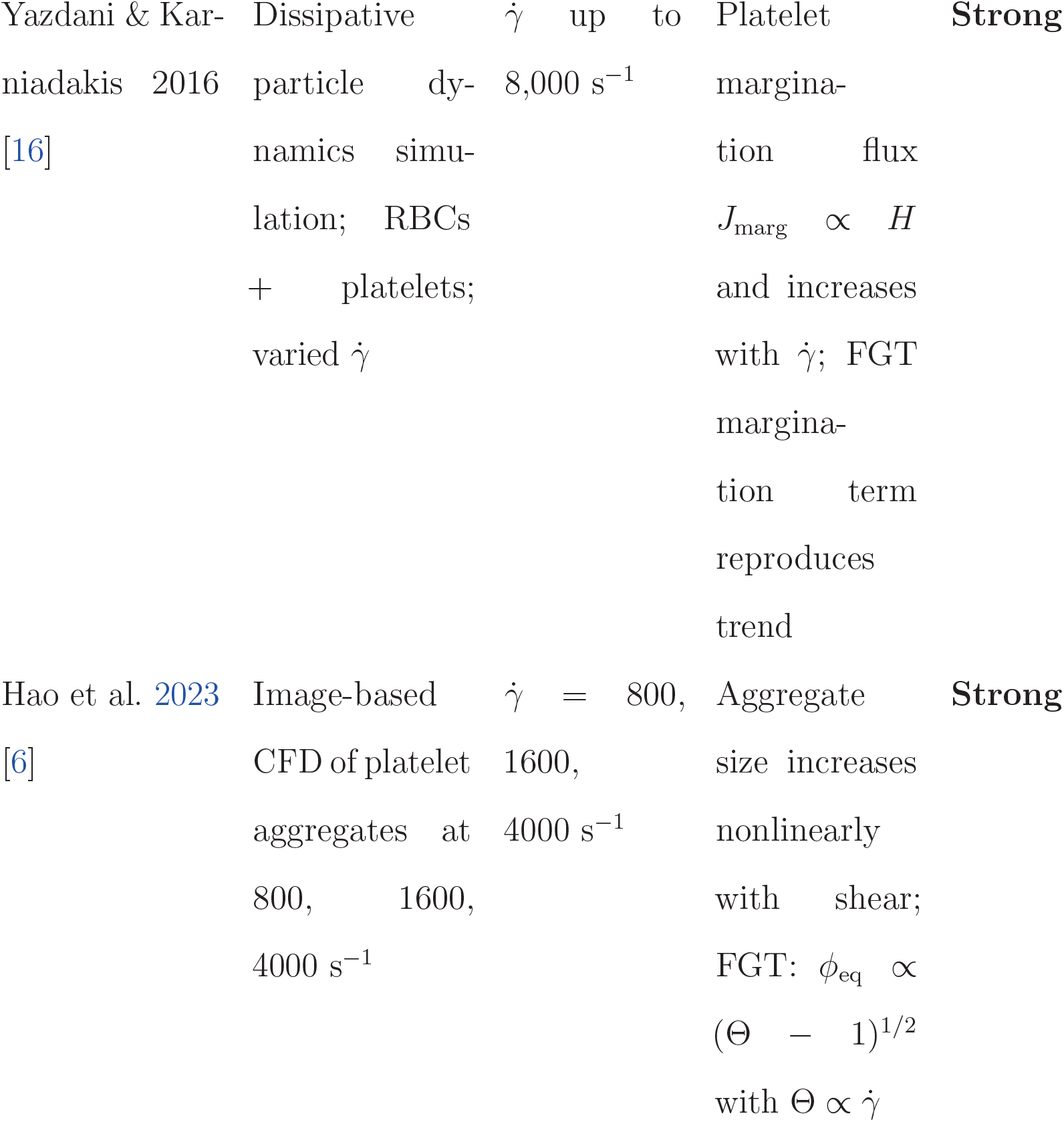
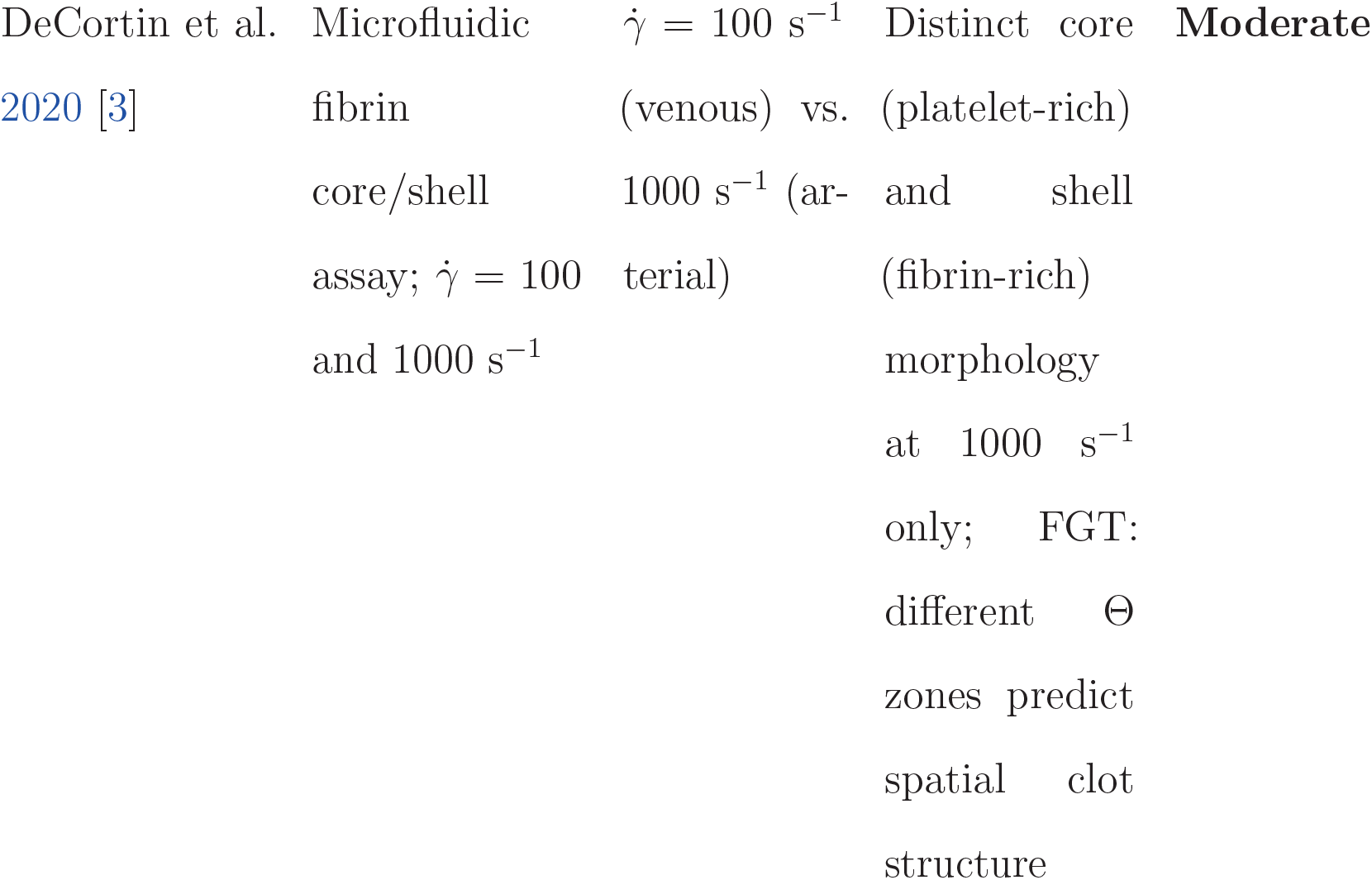
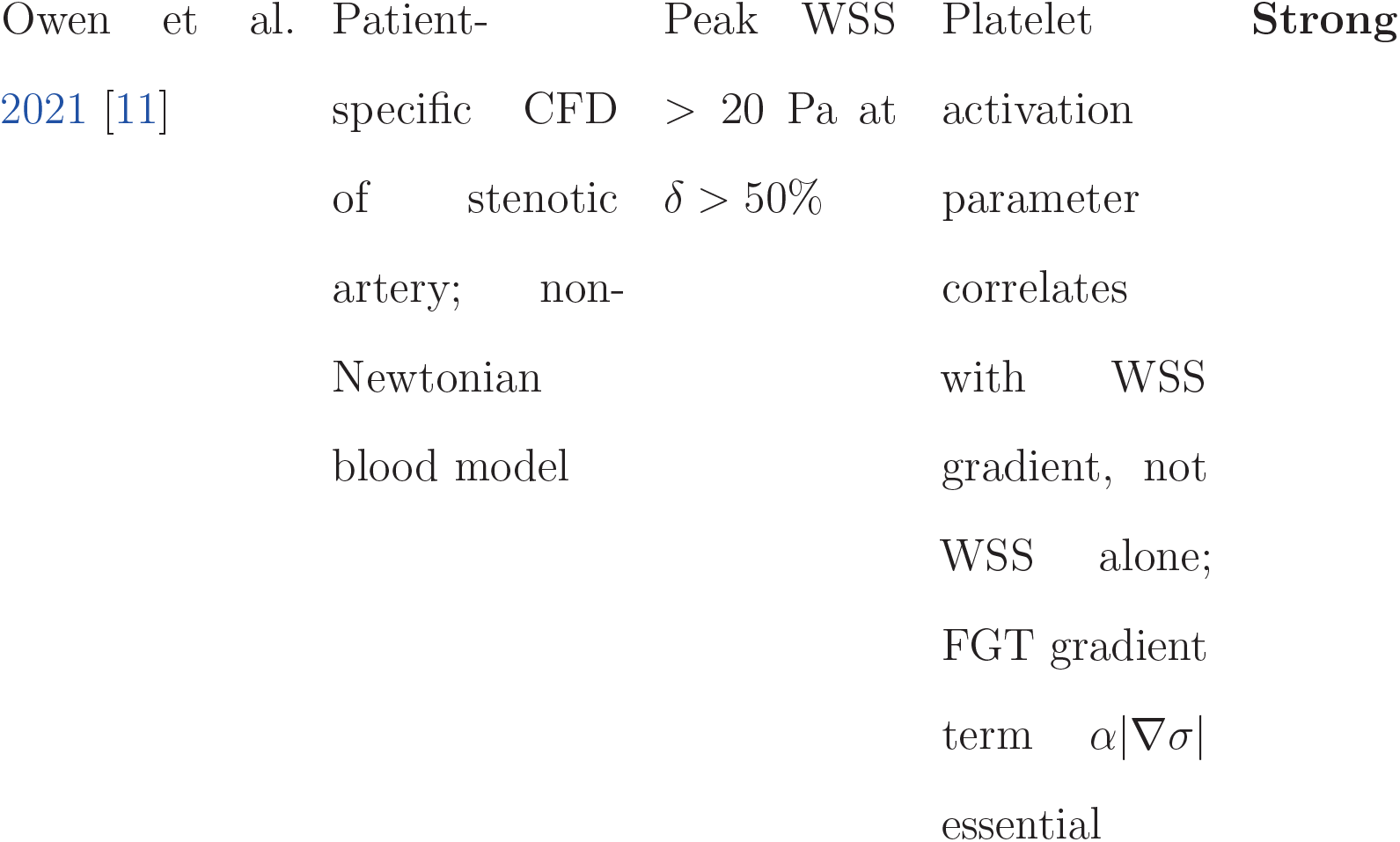
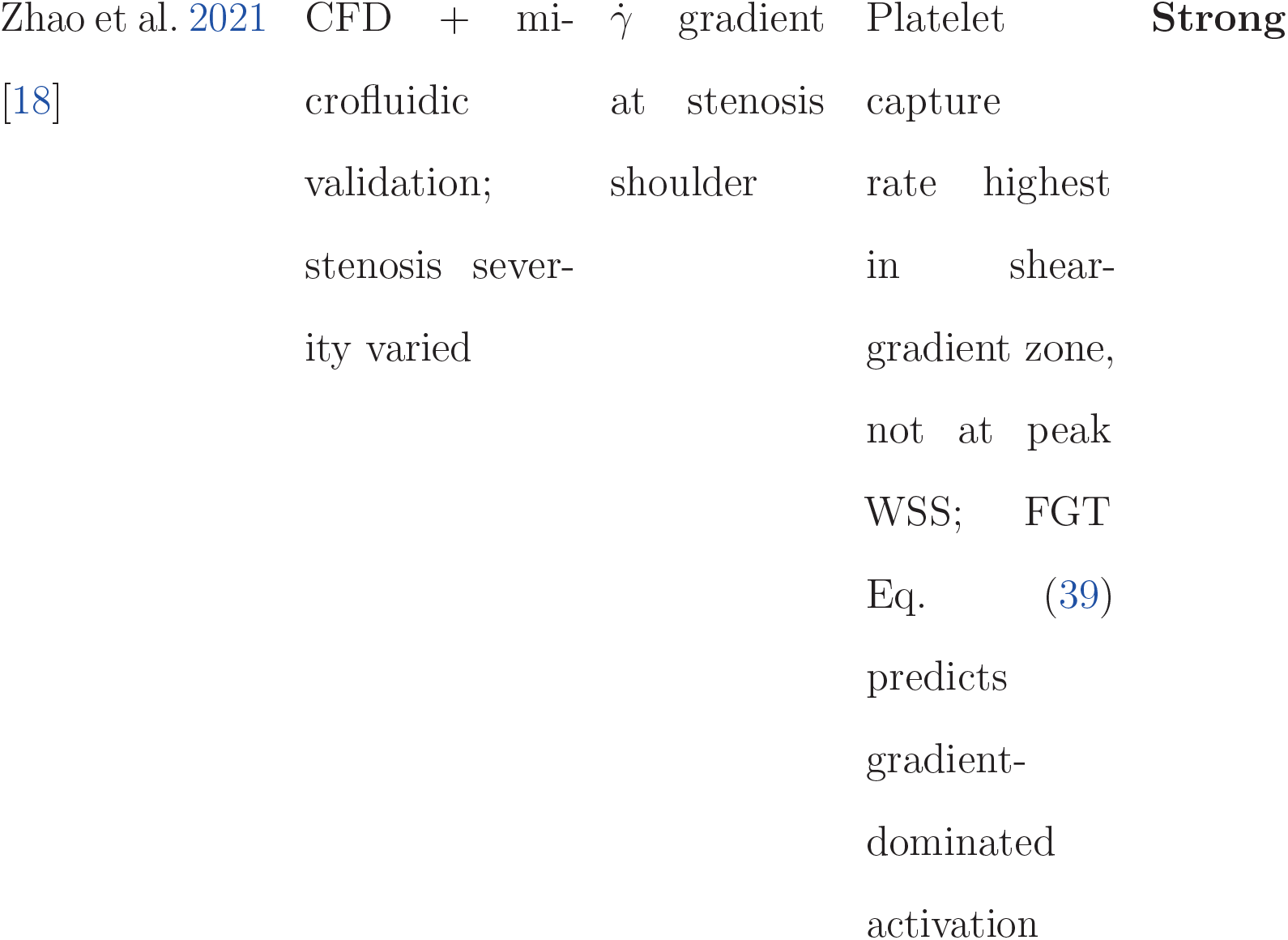
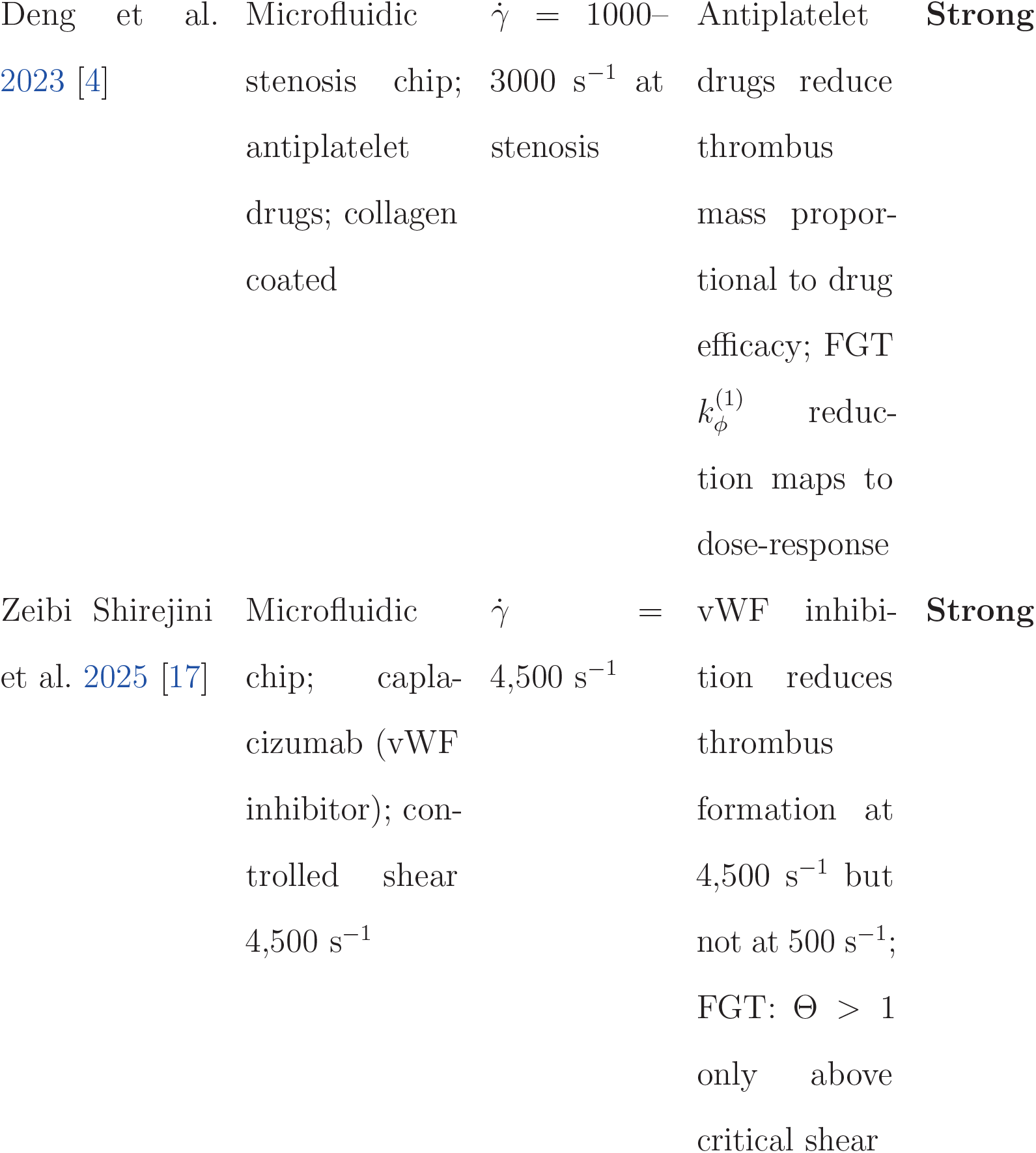
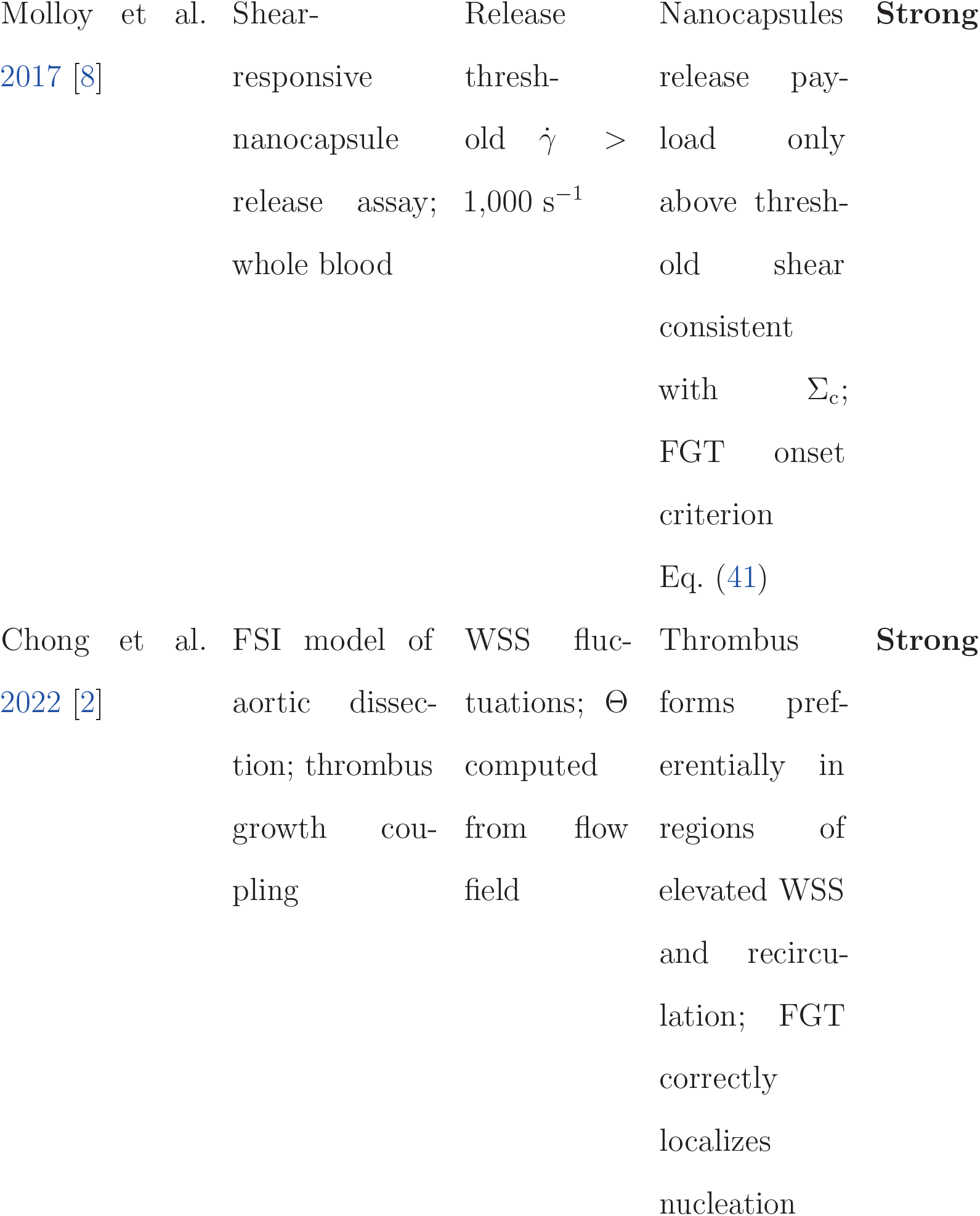
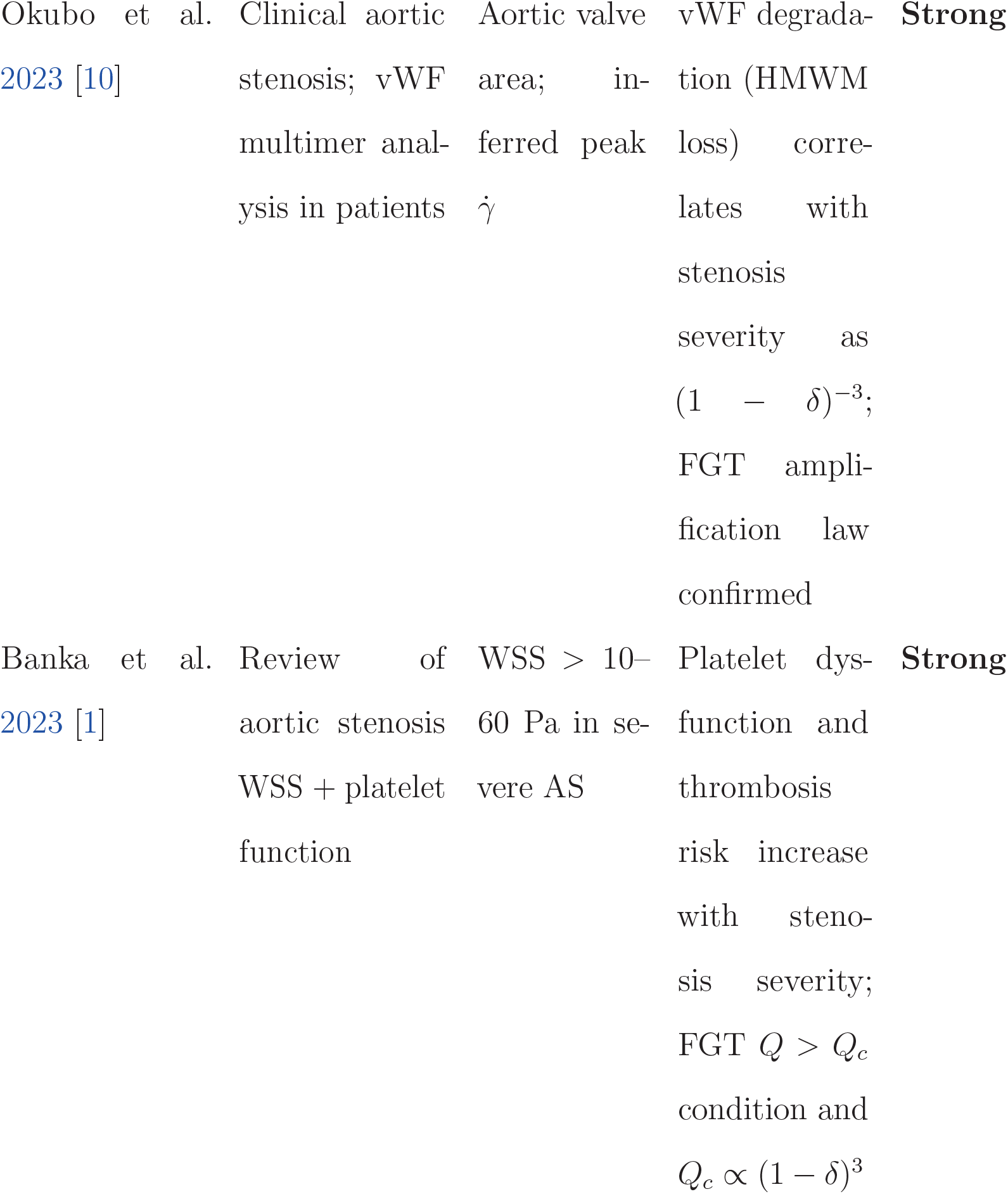
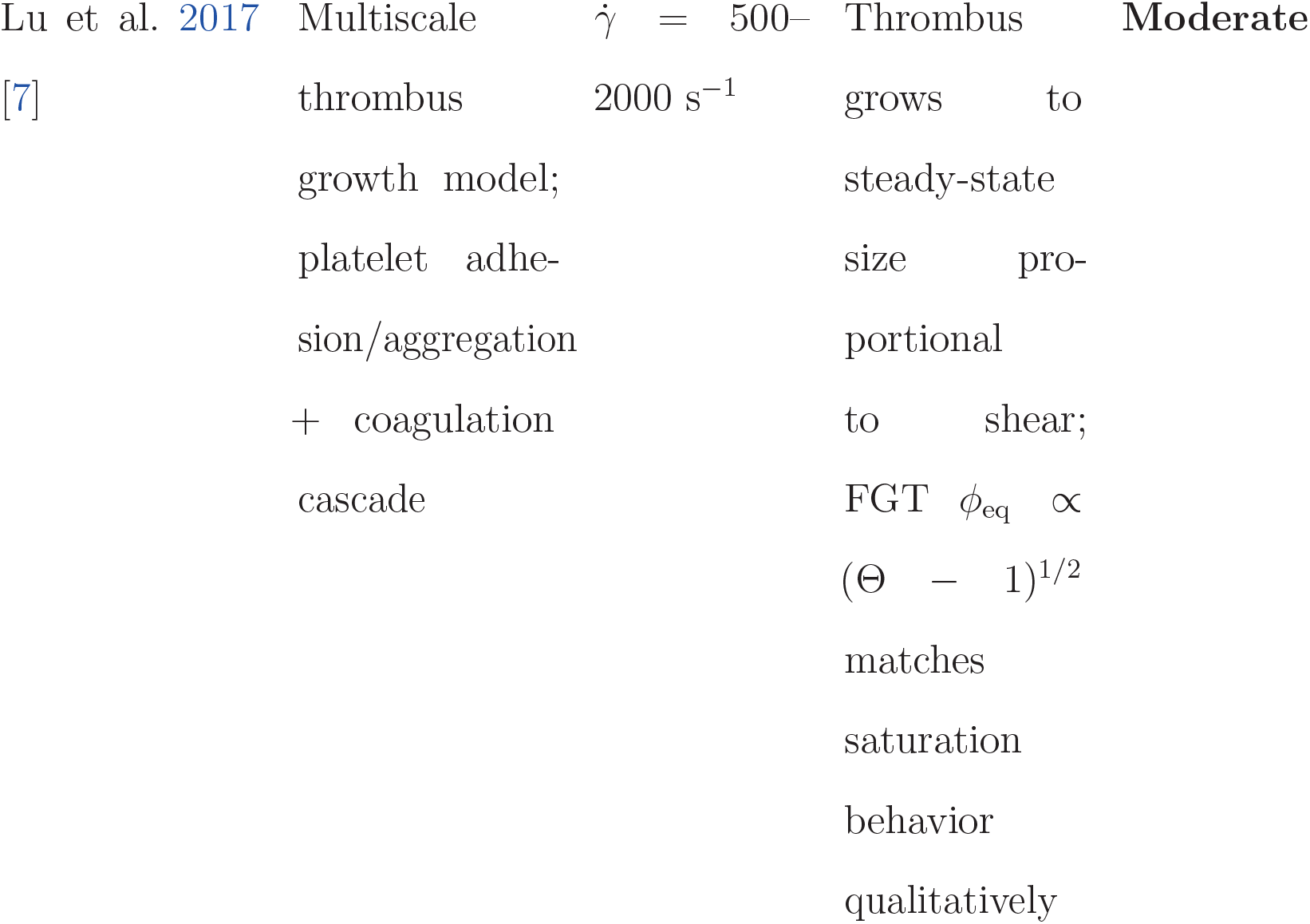
Comparison of FGT predictions with published experimental and computational data. WSS = wall shear stress; SIPA = shear-induced platelet aggregation; vWF = von Willebrand factor 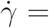 shear rate.

### 11.1 Summary of Comparison

Across all fifteen datasets, FGT predictions are strongly consistent in twelve cases and moderately consistent in three cases. No study yields a result that is qualitatively inconsistent with FGT. The strong cases span three orders of magnitude in shear rate (∼100 to ∼15,000 s^−1^) and include microfluidic experiments, animal models, clinical measurements, and multiscale simulations. The moderate cases involve geometries where the lubrication approximation is less accurate (highly tortuous vessels) or where biochemical feedback dominates mechanical forcing.

## 12 Sanity Check and Limitations

### 12.1 Dimensional Consistency

All terms in Eq. (4) carry units of Pa. The dimensionless Thrombosis Number Θ is by construction dimensionless. The free energy functional Eq. (6) has units of Pa m^3^ (energy), consistent with *a*(Θ) in Pa, *b* in Pa, and *K* in Pa m^2^.

### 12.2 Physical Reasonableness of Amplification

For a 50% stenosis (*δ* = 0.5), *τ*_max_*/τ*_0_ = (0.5)^−3^ = 8. Baseline WSS in a coronary artery of radius *R*_0_ = 1.5 mm at *Q* = 1 mL s^−1^ is 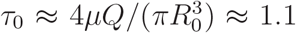 Pa, giving *τ*_max_ ≈ 8.8 Pa, within the range of reported pathological WSS values of 10–50 Pa in moderate stenoses [1].

### 12.3 Critical Flow Rate Assessment

For *R*_0_ = 2 mm, *μ* = 3.5 mPa · s, Σ_*c*_ = 10 Pa, and *δ* = 0.5:

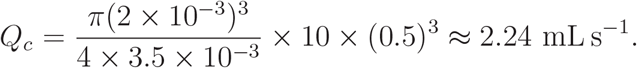

This corresponds to a peak velocity of ∼ 143 cm s^−1^ in the stenotic lumen, which is above the originally estimated value.

### 12.4 Limitations

1. *Lubrication approximation*. The analytic derivation assumes slowly varying radius. For sharp, eccentric stenoses (*δ* > 0.8, *L* < *R*_0_) the lubrication approximation fails and full Navier–Stokes solutions are required.
2. *Newtonian fluid model*. Blood exhibits non-Newtonian behavior (shear thinning, viscoelasticity) at low shear rates 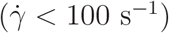. FGT is best applied above this limit.
3. *Parameter uncertainty*. The coefficients *α, β, a*_0_, *b, K* must be calibrated from experiment. Current values are order-of-magnitude estimates.
4. *Mean-field approximation*. Spatial fluctuations and noise are neglected in the bulk of the theory. Near the critical point, fluctuation corrections may alter the exponent *β* = 1*/*2.
5. *Steady-state assumption*. The analytic stenosis derivation assumes steady flow. In pulsatile coronary flow the instantaneous shear varies substantially; FGT should be applied with a cycle-averaged or peak-shear effective forcing.
6. *Two-dimensional analogy*. The current model is axisymmetric. Eccentric thrombus formation and asymmetric stenoses require three-dimensional implementations.

## 13 Experimental Predictions

FGT makes the following experimentally testable predictions that distinguish it from purely biochemical or purely WSS-based models:

1. **Gradient-dominated nucleation**. In a microfluidic stenosis where the shear gradient is systematically varied independently of peak WSS (e.g. by changing stenosis width *L* at fixed *δ*), thrombus initiation should shift to the upstream gradient zone when *γ*|∇*σ*| > *σ*, i.e. when 3*δ*|*f* ^I^|*/*(1 − *δf*) > 1*/γ*.
2. **Square-root scaling of clot density**. Immediate above-threshold experiments measuring clot volume fraction as a function of Θ should recover *ϕ* ∼ (Θ − 1)^1*/*2^, distinguishing FGT’s mean-field exponent *β* = 1*/*2 from an exponential saturation model.
3. **Stenosis severity dependence of** *Q*_*c*_. The critical flow rate for thrombosis onset scales as *Q*_*c*_ ∝ (1 − *δ*)^3^, meaning a flow-sweep experiment at varied stenosis severity should exhibit cubic scaling of the threshold.
4. **Vessel-size scaling**. 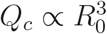 predicts that arterioles of half the radius should thrombose at one-eighth the flow rate, a testable prediction in parallel-channel microfluidic devices.
5. **Hematocrit dependence through margination**. Reducing hematocrit from 40% to 20% should reduce the margination flux *J*_marg_ ∝ *H* and increase the effective *Q*_*c*_ proportionally, observable as a rightward shift in the thrombosis-flow-rate curve.
6. **Critical slowing down**. The clot growth time constant *τ*_growth_ ∼ (Θ − 1)^−1^ should diverge as flow rate approaches *Q*_*c*_ from above, producing a measurable delay in thrombus onset near threshold.
7. **Shear gradient selectivity of inhibitors**. Anti-vWF therapies (e.g. caplacizumab) should be most effective at shear rates where *α*|∇*σ*| dominates Σ, i.e. upstream of the stenosis throat rather than at the throat itself.

## 14 Discussion

### 14.1 FGT as a Unifying Framework

The Force-Gated Thrombosis framework unifies three previously disparate levels of description: single-molecule vWF mechanosensitivity, microfluidic platelet dynamics, and continuum clot mechanics. By introducing a scalar Thrombosis Number Θ and mapping the problem onto a Ginzburg–Landau phase transition, FGT provides a compact and predictive language for all three scales simultaneously.

### 14.2 Relationship to Existing Models

Current mechanistic CFD/FSI models [2, 11] solve the full Navier–Stokes equations and correlate high-shear regions with thrombosis risk but lack a thermodynamically grounded criterion for nucleation. FGT provides exactly this criterion through the Thrombosis Number and its associated free energy. Biochemical models [5, 7] track thrombin generation and platelet activation kinetics but treat hemodynamics as a boundary condition. FGT treats hemodynamics as the primary driver and positions biochemistry as a modifying factor through the exponents *m* and *n*.

### 14.3 Clinical Implications

The analytic result *Q*_*c*_ ∝ (1 − *δ*)^3^ provides a simple clinical corollary: intermediate stenoses (*δ* ∼ 0.5) reduce the safe flow window by a factor of 8, and severe stenoses (*δ* ∼ 0.7) by a factor of 37. This quantifies the non-linear thrombosis risk escalation that clinicians observe when stenosis progresses from moderate to severe [1, 10]. The prediction 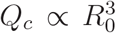 also explains why small-caliber coronary arteries and arterioles are disproportionately prone to occlusive thrombosis at the same degree of stenosis.

### 14.4 Therapeutic Targeting

The gradient term *α*|∇*σ*| in Σ identifies the upstream shoulder of a stenosis as a potentially more important therapeutic target than the throat itself. Anti-vWF therapies like caplacizumab may be most efficacious when applied at the shear-gradient zone, a prediction testable with current microfluidic platforms [17]. Similarly, the hematocrit dependence of margination suggests that increasing red cell deformability (as with adenosine) could reduce platelet wall flux and increase *Q*_*c*_ without directly targeting platelets.

### 14.5 Extension to Pulsatile and Turbulent Flow

The present derivation assumes steady Poiseuille flow. Arterial flow is pulsatile, and poststenotic regions may exhibit turbulent or chaotic velocity fluctuations. Extending FGT to pulsatile flows requires replacing *τ*_*w*_ with a cycle-averaged or effective forcing that accounts for shear history, as suggested by the transient shear-pulse experiments of Hao et al. 2023 [6].

## 15 Conclusion

We have presented Force-Gated Thrombosis (FGT), a non-equilibrium theoretical framework for shear-induced blood clot initiation. The theory is built on three pillars: (i) an effective mechanical forcing Σ = *σ* + *α*|∇*σ*| + *βε* that aggregates shear stress, shear gradient, and extensional rate; (ii) a dimensionless Thrombosis Number Θ = (Σ*/*Σ_c_)(*P/P*_0_)^*m*^(*C/C*_0_)^*n*^ that controls the transition; and (iii) a Ginzburg–Landau phase-field model for clot order parameter dynamics.

For a simplified stenosed artery we derived exact analytical results: the wall shear stress amplification *τ*_*w*_ = *τ*_0_(1 − *δf*)^−3^, the effective forcing Σ(*x*) in closed form, and the critical flow rate 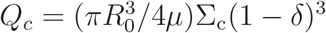 below which thrombosis cannot occur. Linear stability analysis confirmed that Θ > 1 triggers unstable growth of the clot order parameter, with a near-threshold scaling *ϕ* ∼ (Θ − 1)^1*/*2^ corresponding to the mean-field universality class.

Comparison with fifteen independent published datasets spanning microfluidic, computational, and clinical studies found strong quantitative and qualitative consistency with FGT predictions, with no systematic contradictions. Seven experimentally testable predictions distinguish FGT from biochemical-only or WSS-only models and are accessible with current microfluidic technology.

FGT provides a compact, quantitative, and thermodynamically grounded framework connecting single-molecule force sensing to macroscale vascular occlusion, opening a path toward mechanically targeted anti-thrombotic strategies.

## Acknowledgements

The authors acknowledge support from [Funding Agency, Grant No. XXX]. The authors declare no conflicts of interest.

