## Supplementary Information for "Force-Gated Thrombosis (FGT): A Non-Equilibrium Mechanical Theory of Shear-Induced Blood Clot Initiation"

### Supplymentary Information

#### 9A Analytic Derivation for a Rectangular Microfluidic Stenosis Channel

*Supplement to Section 9 of the FGT Theory Paper*

The analytic stenosis derivation in Section 9 assumes a circular cylindrical vessel. In microfluidic experiments, channels are typically fabricated by soft lithography (PDMS on glass), which produces rectangular cross-sections with a fixed height h determined by photoresist thickness. The stenosis is created by laterally narrowing the channel width w(x) while the height remains constant. This geometry fundamentally changes the shear amplification at the stenosis and modifies the critical flow rate scaling law.

##### 9A.1 Geometry of a Rectangular Stenosed Channel

Consider a rectangular microfluidic channel with constant height h and spatially varying width w(x). The width profile is

$$w(x) = w_0(1 - \delta f(x)) \tag{9A.1}$$

where $w_0$ is the width of the upstream channel, $\delta$ is the severity of the stenosis ($0 < \delta < 1$), and $f(x)$ is the shape function of the stenosis, identical to Eq. (26). The minimum width at the throat is

$$w_min = w_0(1 - \delta) \tag{9A.2}$$

The key geometric distinction from Section 9 is that the cross-sectional area scales as $A(x) = w(x) \cdot h$, which is linear in w, whereas for a circular pipe $A = \pi R^2$ is quadratic in R. This difference propagates through the entire derivation.

##### 9A.2 Wall Shear Stress in a Rectangular Channel

For a rectangular channel with $w \gg h$ (the shallow-channel or Hele–Shaw limit), the flow is approximately parabolic across the height, and the wall shear stress on the top and bottom walls is

$$\tau_w(x) = \frac{6\mu Q}{w(x)h^2}. \tag{9A.3}$$

Compare Eq. (32) for a circular pipe,

$$\tau_w = \frac{4\mu Q}{\pi R^3}.$$

The critical difference is the exponent: for a fixed-height rectangular channel,

$$\tau_w \propto \frac{1}{w},$$

whereas for a circular pipe,

$$\tau_w \propto \frac{1}{R^3}.$$

Thus, the cubic dependence on the constricted radius applies to circular or geometrically similar stenoses, but not directly to a fixed-height rectangular microchannel in which only the lateral width changes [1, 2]..

**Finite-aspect-ratio correction.** Eq. (9A.3) is exact in the parallel-plate limit $w/h \to \infty$. For finite aspect ratios, the top–bottom wall shear stress can be written more generally as

$$\tau_w(x) = \frac{6\mu Q}{w(x)h^2}\,\Phi\left(\frac{w(x)}{h}\right),$$

where $\Phi(w/h) \to 1$ as $w/h \to \infty$. For sufficiently large aspect ratios, for example $w/h \gtrsim 3$, the correction is small and Eq. (9A.3) is adequate as a leading-order design estimate.

To make this correction explicit, we use the standard leading finite-aspect-ratio approximation for a straight rectangular duct, in which the hydraulic resistance contains the factor [3, 4, 2].

$$1 - 0.63\frac{h}{w}, \qquad w \geq h.$$

This approximation is commonly used for laminar flow in rectangular microchannels and follows from the rectangular-duct series solution. It gives the scaling

$$\tau_w(x) \sim \frac{1}{w(x)\,(1 - 0.63h/w(x))} = \frac{1}{w(x) - 0.63h}. \qquad (9A.4)$$

This expression should be interpreted as a finite-aspect-ratio refinement of the transparent $1/w(x)$ scaling, not as a replacement for the CFD wall-traction calculation. It shows that, as $w(x)/h$ approaches order unity, the wall shear stress increases more strongly than predicted by the simple parallel-plate estimate.

Using the previously defined stenosis geometry,

$$w(x) = w_0[1 - \delta f(x)],$$

the normalized finite-aspect-ratio-corrected wall shear stress becomes

$$\frac{\tau_w(x)}{\tau_0} = \frac{1 - 0.63h/w_0}{(1 - \delta f(x)) - 0.63h/w_0}. \qquad (9A.5)$$

Differentiating Eq. (9A.5) with respect to $x$ gives

$$\frac{1}{\tau_0}\frac{d\tau_w}{dx} = \delta f'(x)\frac{1 - 0.63h/w_0}{\left[(1 - \delta f(x)) - 0.63h/w_0\right]^2}. \qquad (9A.6)$$

Taking the magnitude,

$$\boxed{\frac{1}{\tau_0}\left|\frac{d\tau_w}{dx}\right| = \delta|f'(x)|\frac{1 - 0.63h/w_0}{\left[(1 - \delta f(x)) - 0.63h/w_0\right]^2}} \qquad (9A.7)$$

In the parallel-plate limit $h/w_0 \to 0$, Eq. (9A.7) reduces to the leading-order rectangular result,

$$\frac{1}{\tau_0}\left|\frac{d\tau_w}{dx}\right| = \delta|f'(x)|(1 - \delta f(x))^{-2}. \qquad (9A.8)$$

This limiting expression captures the dominant $1/w(x)$ dependence, while Eq. (9A.7) quantifies the finite-aspect-ratio deviation when $w/h$ is not large.

For the present $300\,\mu\text{m} \times 100\,\mu\text{m}$ channel, the throat aspect ratios are

$$w_{\text{throat}}/h = 2.1,\ 1.5,\ \text{and } 0.9$$

For the present $300\,\mu\text{m} \times 100\,\mu\text{m}$ channel, the throat aspect ratios are

$$w_{\text{throat}}/h = 2.1,\ 1.5,\ \text{and } 0.9$$

for the 210, 150, and 90 $\mu$m throat widths, respectively. Therefore, the finite-aspect-ratio correction is expected to be most reliable for the mild and intermediate stenosis cases where $w_{\text{throat}}/h \geq 1.5$, corresponding to $\delta \leq 0.5$ in this device. For more severe stenosis, where $w_{\text{throat}}/h$ drops below unity, the parallel-plate-based framework breaks down and CFD is required for quantitative WSSG extraction.

This behaviour is quantified in Tables 1 and 2. For the 30% and 50% stenosis cases, the finite-aspect-ratio correction improves the CFD/formula peak ratio to approximately 1.10, indicating good peak-level agreement. In contrast, for the 70% stenosis case, the corrected analytical expression overcorrects the WSSG and gives a CFD/corrected-formula peak ratio of approximately 0.61. This indicates that the finite-aspect-ratio correction should not be used as a quantitative predictor once $w_{\text{throat}}/h < 1$, and that CFD wall-traction extraction is required in this regime.

Table 1: Geometry and aspect-ratio conditions for the three rectangular stenosis cases. The finite-aspect-ratio correction is expected to be most reliable for $w_{\text{throat}}/h \geq 1.5$.

| Stenosis | $w_{\text{throat}}$ ($\mu$m) | $h$ ($\mu$m) | $w_{\text{throat}}/h$ |
|---|---|---|---|
| 30% | 210 | 100 | 2.1 |
| 50% | 150 | 100 | 1.5 |
| 70% | 90 | 100 | 0.9 |

Table 2: Comparison between CFD-derived peak WSSG and analytical estimates. Peak ratios are defined as $W_{\text{CFD,peak}}/W_{\text{formula,peak}}$.

| Stenosis | CFD peak (1/m) | Simple peak (1/m) | CFD/simple | Corrected peak (1/m) | CFD/corrected |
|---|---|---|---|---|---|
| 30% | 741.19 | 462.80 | 1.60 | 676.45 | 1.10 |
| 50% | 2447.57 | 1165.81 | 2.10 | 2217.31 | 1.10 |
| 70% | 9876.21 | 3231.04 | 3.06 | 16300.36 | 0.61 |

Together, these results support a two-level interpretation of the rectangular-channel model. The leading-order expression captures the physically transparent $1/w(x)$ scaling, while the finite-aspect-ratio correction quantifies the expected deviation when $w/h$ is not large. However, when the stenosis becomes sufficiently severe that $w_{\text{throat}}/h < 1$, the analytical correction no longer provides a reliable quantitative prediction and instead overcorrects the WSSG. Accordingly, the analytical expressions are used to interpret scaling and WSSG localization, whereas the CFD wall-traction values are used as the quantitative ground truth for subsequent mechanical and thrombosis-threshold analyses.

#### 9A.3 Shear Amplification by Stenosis

Substituting Eq. (9A.1) into Eq. (9A.3):

$$\tau_w(x) = \frac{6\mu Q}{w(x)h^2} \cdot (1 - \delta f(x))^{-1} \tag{9A.9}$$

Defining the baseline wall shear stress $\tau_0 = 6\mu Q/(w_0 h^2)$:

$$\tau_w(x) = \tau_0 \cdot (1 - \delta f(x))^{-1} \tag{9A.10}$$

At the stenosis throat (x = 0, f = 1):

$$\tau_{\max} = \tau_0 \cdot (1 - \delta)^{-1} \tag{9A.11}$$

Compare the circular result Eq. (35): $\tau_{\max} = \tau_0(1-\delta)^{-3}$. The amplification factor is $(1-\delta)^{-1}$ rather than $(1-\delta)^{-3}$, reflecting the reduced geometric confinement when only one dimension narrows.

#### Amplification Comparison

| $\delta$ | Rectangular $(1-\delta)^{-1}$ | Circular $(1-\delta)^{-3}$ | Ratio |
|---|---|---|---|
| 0.3 | ×1.43 | ×2.92 | 0.49 |
| 0.5 | ×2.00 | ×8.00 | 0.25 |
| 0.7 | ×3.33 | ×37.04 | 0.09 |

*Table 9A.1: Shear amplification at different stenosis severities for rectangular and circular geometries. At severe stenosis $\delta = 0.7$, the rectangular geometry gives only a ×3.33 amplification, compared with ×37.04 for a circular pipe.*

#### 9A.4 Shear Gradient

Differentiating Eq. (9A.10):

$$d_{\tau w}/dx = \tau_0 \delta f'(x)(1 - \delta f(x))^{-2} \tag{9A.12}$$

Compare Eq. (36): $\frac{d\tau_w}{dx} = 3\tau_0\delta f'(x)(1-\delta f(x))^{-4}$. The rectangular gradient has a prefactor of 1 rather than 3, and an exponent of −2 rather than −4, reflecting the weaker geometric focusing. The magnitude is:

$$|\nabla\tau| = \tau_0\delta\,|f'(x)|\,(1-\delta f(x))^{-2}\,. \tag{9A.13}$$

#### 9A.5 Effective Mechanical Forcing

Following Eq. (38) and defining $\gamma = \alpha + \beta\kappa$:

$$\Sigma(x) = \tau_0 \left(1 - \delta f(x)\right)^{-1} + \gamma\tau_0\delta \left|f'(x)\right| \left(1 - \delta f(x)\right)^{-2}. \tag{9A.14}$$

Compare the circular result Eq. (39): $\Sigma(x) = \tau_0 \left(1 - \delta f(x)\right)^{-3} + 3\gamma\tau_0\delta \left|f'(x)\right| \left(1 - \delta f(x)\right)^{-4}$.

#### 9A.6 Thrombosis Number and Onset Criterion

Setting $P = P_0$, $C = C_0$ (or equivalently $m = n = 0$ for a pure mechanical experiment):

$$\Theta(x) = \frac{\Sigma(x)}{\Sigma_c} = \frac{1}{\Sigma_c}\left[\tau_0 \left(1 - \delta f(x)\right)^{-1} + \gamma\tau_0\delta \left|f'(x)\right| \left(1 - \delta f(x)\right)^{-2}\right]. \tag{9A.15}$$

At the throat ($f = 1,\ f' = 0$), the gradient term vanishes:

$$\Sigma_{throat} = \tau_0(1 - \delta)^{-1} \tag{9A.16}$$

The thrombosis onset condition $\Theta > 1$ becomes:

$$\tau_0(1 - \delta)^{-1} > \Sigma_c \tag{9A.17}$$

#### 9A.7 Critical Flow Rate for Rectangular Channels

Substituting $\tau_0 = 6\mu Q/(w_0 h^2)$ into Eq. (9A.17) and solving for Q:

$$Q_c^{\text{rect}} = \frac{w_0 h^2}{6\mu}\,\Sigma_c\,(1 - \delta). \tag{9A.18}$$

Compare the circular result Eq. (44): $Q_c = \frac{\pi R_0^3}{4\mu}\Sigma_c(1 - \delta)^3$.

The fundamental result: **for a constant-height rectangular channel, the critical flow rate scales linearly with stenosis severity, $Q_c \propto (1 - \delta)$, rather than the cubic scaling $Q_c \propto (1 - \delta)^3$ of circular pipes.**

#### 9A.8 Physical Interpretation

The weakened scaling has a clear physical origin. In a circular pipe, a 50% stenosis ($\delta = 0.5$) reduces the radius by half, but the cross-sectional area by a factor of 4 (since $A \propto R^2$). The velocity must increase by $\times 4$ to maintain flow rate, and the velocity gradient at the wall increases even further ($\propto 1/R^3$) due to the tighter curvature. In a constant-height rectangular channel, the same 50% stenosis reduces only the width, cutting the area by half. The velocity increases by $\times 2$, and the wall shear stress increases by only $\times 2 (\propto 1/w)$. The height dimension, being constant, provides no additional geometric focusing. This has a direct clinical analogue: in vivo, atherosclerotic stenoses constrict arteries circumferentially, producing the cubic scaling. Microfluidic models that narrow only one dimension systematically underestimate the shear amplification for a given stenosis severity.

#### 9A.9 Unified Scaling Law

Both cases are subsumed by a general scaling law. If the wall shear stress scales as $\tau_w \propto Q/\ell_p$, where $\ell$ is the characteristic length that narrows and p is the geometry-dependent exponent, then:

$$Q_c \propto (1 - \delta)p \tag{9A.19}$$

| Geometry | $\tau_w$ scalling | $Q_c$ exponent p |
|---|---|---|
| Circular pipe (R narrows) | $\tau \propto Q/R^3$ | 3 |
| Square channel (a narrows) | $\tau \propto Q/a^3$ | 3 |
| Rectangular, w and h both narrow $\tau$ | $\tau \propto Q/(wh^2)$ | 3 |
| Rectangular, only w narrows ($h = const$) | $\tau \propto Q/w$ | 1 |
| Slit channel (only h narrows, $w = const$) | $\tau \propto Q/h^2$ | 2 |

*Table 9A.2: Unified scaling law for different channel geometries. The exponent p equals the power of the narrowing dimension in the wall shear stress formula.*

#### 9A.10 Near-Threshold Scaling

The near-threshold behaviour of the order parameter is unchanged:

$$\varphi \sim \left(\frac{Q}{Q_c} - 1\right)^{1/2}. \tag{9A.20}$$

The mean-field exponent $\beta = 1/2$ is a property of the Ginzburg–Landau free energy, not of the channel geometry. What changes is the value of $Q_c$ itself, and therefore the absolute flow rate at which the transition occurs

#### 9A.11 Numerical Example

For a representative microfluidic device with $w_0 = 300\ \mu$m, $h = 100\ \mu$m, $\mu = 3.5$ mPa · s (whole blood), $\Sigma_c = 15$ Pa, and $\delta = 0.5$,

$$Q_c = \frac{(300 \times 10^{-6})(100 \times 10^{-6})^2 \times 15 \times 0.5}{6 \times 3.5 \times 10^{-3}}$$

$$= \frac{2.25 \times 10^{-11}}{2.1 \times 10^{-2}} = 1.07 \times 10^{-9}\ \text{m}^3/\text{s}$$

$$\approx 64\ \mu\text{L/min}.$$

For water ($\mu = 1$ mPa · s), the equivalent flow rate is

$$Q_{c,\text{water}} = \left(\frac{\mu_{\text{blood}}}{\mu_{\text{water}}}\right) Q_c = 3.5 \times 64 \approx 224\ \mu\text{L/min},$$

confirming that the transition is accessible with standard syringe pumps.

#### 9A.12 Summary of Rectangular Channel Results

This derivation yields: (i) a leading-order wall shear stress amplification law

$$\tau_w \propto (1-\delta)^{-1}$$

for fixed-height rectangular channels, in contrast to the circular-pipe scaling

$$\tau_w \propto (1-\delta)^{-3};$$

(ii) a finite-aspect-ratio refinement showing that the transparent $1/w(x)$ scaling can be corrected as

$$\tau_w(x) \sim \frac{1}{w(x) - 0.63h},$$

with improved CFD/formula peak agreement for $w_{\text{throat}}/h \geq 1.5$, but loss of quantitative reliability once $w_{\text{throat}}/h < 1$;(iii) an effective forcing $\Sigma(x)$ in closed form, with the WSSG contribution determined by the local narrowing function $f(x)$ and wall slope $f'(x)$; (iv) a critical flow rate

$$Q_c = \frac{w_0 h^2}{6\mu}\Sigma_c(1-\delta),$$

giving the leading-order linear stenosis scaling

$$Q_c \propto (1-\delta)$$

for fixed-height rectangular channels; (v) a unified scaling law

$$Q_c \propto (1-\delta)^p,$$

where $p = 1$ for fixed-height shallow rectangular stenoses and $p = 3$ for circular or geometrically similar stenoses; and (vi) confirmation that the near-threshold scaling

$$\phi \sim (Q/Q_c - 1)^{1/2}$$

is geometry-independent once the appropriate geometry-dependent $Q_c$ is used.

Together, these results show that rectangular microfluidic stenosis models should not be interpreted using circular-pipe stenosis scaling without correction. The analytical expressions provide scaling insight and clarify how stenosis severity, wall slope, and channel geometry affect WSSG, while CFD wall-traction values provide the geometry-specific quantitative WSSG used for subsequent mechanical and thrombosis-threshold analysis. Quantitative translation between fixed-height microfluidic models and circular or in vivo vessels therefore requires accounting for the geometry-dependent exponent $p$, finite-aspect-ratio effects, and, where necessary, CFD-derived wall-traction values.

### 10A Ghost Particle Velocimetry

*Supplement to Section 10 of the FGT Theory Paper*

Ghost Particle Velocimetry (GPV) was used to reconstruct two-dimensional flow velocity fields in the micro/milli-fluidic devices. GPV is a particle-based velocimetry technique that relies on the cross-correlation of speckle patterns generated by nanoparticles dispersed in the working fluid. In contrast to conventional micro-particle image velocimetry (µPIV), GPV employs tracer particles with diameters below the optical diffraction limit, enabling flow-field measurements with minimal disturbance while using a standard bright-field microscopy configuration. The method was implemented following the principles reported by Riccomi et al. [5].

The GPV experiments were performed using an inverted optical microscope equipped with a white LED illumination source. To generate the speckle pattern required for GPV analysis, the numerical aperture of the condenser lens, $NA_c$, was reduced by partially closing the condenser aperture diaphragm to approximately 0.15–0.20. This adjustment produces partially coherent illumination within a confined measurement volume, allowing speckle formation from the dispersed nanoparticles. The characteristic thickness of this illuminated volume, $\delta$, is related to the illumination wavelength, $\lambda$, and condenser numerical aperture according to

$$\delta = \frac{\lambda}{NA_c}. \tag{9A.21}$$

Polystyrene nanoparticles with a nominal diameter of 200 nm were used as flow tracers. Because their diameter is below the diffraction limit of the optical system, individual particles were not directly resolved in the raw images. Instead, the collective scattering from the nanoparticle suspension produced a detectable speckle pattern suitable for velocity-field reconstruction. The particles were suspended in deionized water at a concentration of 0.1% w/w, which provided sufficient optical signal while minimizing particle–particle interactions and avoiding measurable disturbance to the flow. The estimated Stokes number was in the range of $10^{-3}$–$10^{-4}$, indicating negligible particle inertia and confirming that the nanoparticles faithfully followed the fluid motion.

A high-speed camera was mounted directly on the microscope to record time-resolved image sequences of the evolving speckle pattern. The acquisition frame rate was selected according to the imposed flow velocity to ensure an appropriate displacement of the speckle pattern between consecutive frames for cross-correlation analysis. The nanoparticle-seeded fluid was delivered to the device using a syringe pump operating at a constant volumetric flow rate, with microbore tubing used to connect the syringe to the device inlet and to collect the outflow.

Prior to each experiment, the particle suspension was filtered through a 0.45 µm syringe filter to reduce dust and large contaminants. Any remaining impurities observed in the recorded images were removed during post-processing, as their displacement vectors were readily distinguishable from those associated with the speckle pattern.

Image sequences were preprocessed to isolate the dynamic speckle signal from the stationary transmitted-light background. For each experiment, a background image was generated by calculating the median intensity projection of up to 600 frames. This median image was then subtracted from each frame in the sequence using ImageJ [6], thereby removing static optical features and enhancing the fluctuating speckle pattern produced by the moving nanoparticles.

Velocity fields were reconstructed by applying cross-correlation analysis to consecutive image pairs. The preprocessed image sequences were analysed using PIVlab, an open-source MATLAB-based particle image velocimetry routine [7]. A two-pass correlation procedure was used. The first pass employed interrogation windows of $32 \times 32$ pixels, followed by a second pass with reduced interrogation windows of $16 \times 16$ pixels. Fast Fourier Transform window-deformation cross-correlation with linear window deformation was applied, and the correlation step size was set to half of the interrogation-window size to improve spatial resolution. The resulting velocity vectors were exported as comma-separated values files containing the in-plane velocity components at each interrogation point. Custom MATLAB scripts were then used to process the exported data and generate velocity-field contour plots for quantitative analysis and visualization.
